# Creating complete life histories of individual female tsetse (*Glossina* spp) to study the effects of meteorological conditions on fly size in Zimbabwe

**DOI:** 10.64898/2026.03.06.710017

**Authors:** John W. Hargrove, Faikah Bruce, John Van Sickle

## Abstract

Combining novel methodologies with ovarian dissection, we estimated life histories for *ca.* 90,000 individual female *Glossina pallidipes* and *G. m. morsitans* sampled from 1988-1999 in Zimbabwe’s Zambezi Valley. Using temperature-dependent development rates we stepped back through each fly’s life, fixing dates of successive pregnancies, adult emergence, pupal period, pregnancy and oogenesis. This enabled modelling of relationships between wing and egg lengths, and conditions prevailing when these lengths were being determined. Egg lengths increased with maternal wing length, were shorter in primiparous flies but changed little with age thereafter. *G. pallidipes* egg lengths were positively related to NDVI and negatively to temperature (R^2^ = 0.68), for variables averaged over the period of oogenesis for each fly, and then averaged again across weekly cohorts of flies. *G. m. morsitans* mean egg lengths, pooled by month, showed the same pattern (R^2^ = 0.53). Pooled mean wing lengths increased with NDVI and decreased with temperature prevailing while flies were developing in the ovaries and uterus; R^2^= 0.66 (*G. pallidipes*) and R^2^= 0.56 (*G. m. morsitans*). The models – fitted using flies captured after November 1991 – gave good predictions, with no further modeling, for egg and wing lengths of flies captured between September 1988 and November 1991. The models facilitate true predictions of future changes in fly size based on readily available meteorological data, benefiting vector and disease control efforts in predicting likely changes in tsetse population densities and distribution. Selection against small individuals in the hot-dry season is not restricted to teneral mortality continuing for some weeks after emergence. NDVI, measures of wetness and temperature can indirectly impact tsetse size, mortality and population density by affecting vertebrate host density and, thereby, the probability of tsetse locating and feeding on a host. Our methodology impacts numerous areas of vector biology and control.

## Introduction

Adult tsetse flies (*Glossina* spp, Diptera: Glossinidae) are the vectors of human and animal trypanosomiasis (HAT and AAT) in Africa and their field biology has been the subject of intense study since the early 20^th^ century. There have been impressive strides recently in the elimination of the Gambian form of trypanosomiasis and vector control plays an important part in this process (Mahamat et al., 2017; Tirados et al., 2020; Hope et al., 2022). Improved understanding of tsetse biology and population dynamics can thus continue to play a significant role in disease control.

Reproduction in tsetse involves the production, at approximately 9-day intervals, of a single larva that weighs as much or more than its postpartum mother. Once deposited the larva does not feed. Instead, it burrows into loose soil then forms around itself a hard puparial case, within which metamorphosis is completed. The teneral adult fly that emerges from the puparial case, and hardens, has the linear dimensions of the mature adult. The size of the wing, which is commonly used as a measure of fly size, is fully determined by the end of the pupal stage. Thereafter, wing size does not increase with age and only shortens in the adult stage due to wearing at the wing margins. Similarly, the length of an egg is fixed by the time it is ovulated into the uterus (Denlinger & Ma, 1974).

As with all insects, the size of individuals of a given species of tsetse (*Glossina* spp) is constrained within relatively tight limits though individual sizes vary consistently with season. It has been argued that size variation reflects the physical conditions experienced by the individual’s mother while she was bearing the oocytes, egg and larval stages (Jackson, 1948 b, 1953; Bursell & Glasgow, 1960; Glasgow, 1961; Glasgow & Bursell, 1961; Randolph & Rogers, 1986; Rogers, 1991; Dransfield et al., 1989; Hargrove et al., 2019). The central problem that bedevils all such studies was recognised by Glasgow & Bursell (1961) who noted that “The demonstration of a significant correlation should be the beginning, not the end, of such investigations, since it is essential to establish whether the causal chain postulated as linking the variables does in fact exist”; 64 years later no such causal link has yet been established.

Jackson (1953) used wing fray measurements to estimate a mean age of 3-4 weeks for male tsetse collected near Shinyanga in Tanzania (Jackson, 1946, 1948a). Since the pupal period is about 35 days for males at the prevailing mean temperature of 23°C, he expected that the size of male flies of average age 3-4 weeks would be determined approximately 2 months prior to collection, at the time when the fly’s mother was pregnant. He found a negative correlation between mean wing vein length and mean saturation deficit 2 months prior to capture. This correlation was strongest for *G. pallidipes* Austen but was also significant for *G. swynnertoni* Austen and *G. m. morsitans* Westwood. Correlations were progressively reduced as the time lag was reduced to one month and zero months and the correlation was weaker with temperature than with saturation deficit. Note that, whereas all males were assigned the average age of 3-4 weeks, some flies will have been *in utero* for less and some for more time, than the two-month lag period assumed for all flies.

The studies mentioned above all involved the analysis of male tsetse, since females were much more difficult to sample using the handnet method of sampling used at the time. Females are of greater interest, however, and with the advent of odor-baited traps were readily captured. Dransfield *et al*. (1989) analyzed newly emerged female *G. pallidipes* trapped at Nguruman, Kenya, close to the equator. Assuming constant durations of pregnancy and pupal duration throughout the year, they estimated that pregnancy occurred approximately two months prior to sampling. Wing lengths were positively correlated with minimum relative humidity and negatively correlated with maximum temperature, maximum saturation deficit and mean soil temperature. The advantage of this study was that it pinpointed the timing of pregnancy by using flies of known age; its shortcoming was that it used information only from the youngest flies, which generally comprise <7% of flies sampled.

In this paper, we take up the challenge of relating fly sizes to environmental factors. We introduce new methodologies that allow us to estimate entire life histories, for each of 87,000 field-captured female tsetse, and thereby identify the period during which each of a very large number of individuals was developing in its mother’s ovaries and uterus. We then compute averages of meteorological variables over this developmental period for each fly and use correlation and regression to relate these averages to fly size. We included all females for which the chronological age and pupal duration could be estimated and have developed a method for estimating the calendar period when that fly was developing inside its mother, either as an oocyte in the ovaries, or as an egg or larva *in utero*. This has been achieved using subsets of data arising from an 11-year study involving the ovarian dissection of *ca*. 154, 500 *G. pallidipes* and *ca*. 19, 500 *G. m. morsitans*. Since the work was performed in Zimbabwe, relatively far from the equator, seasonal variation in meteorological conditions was much greater than in the East African localities of the studies cited above.

## Methods

### Meteorological measurements

Daily maximum and minimum temperatures were recorded from a mercury thermometer in a Stevenson screen at Rekomitjie; a rain gauge sited next to the screen, likewise, produced daily records of precipitation. From 17 November 1991 – 31 December 1999, hourly mean measurements of shade temperature and relative humidity were also made using an automatic weather station (type WS01, Delta-T devices, Newmarket, UK) sited *ca*. 200 m from the Stevenson screen. Temperature and relative humidity data from this logger were used to calculate the saturation deficit – an index of humidity characterized by the difference between the saturation vapor pressure and the actual vapor pressure of a volume of air. Saturation deficit is thus a pressure, with units conveniently expressed in millibar (mB) and has the property of being proportional to the evaporation capability of the air.

We also estimated the number of hours available for tsetse feeding each day, as a potential explanatory variable. This variable may alter the probability that a female fly finds and feeds on a host on a given day – and may therefore impact her nutrition during pregnancy. Tsetse are generally inactive at night, and at low and high temperatures (Vale, 1971). Accordingly, any hour of a given day was judged suitable for tsetse feeding activity if it lay between the recorded times of sunrise and sunset at Rekomitjie and if the mean temperature for that hour was ≥ 16 and ≤ 32°C.

Daily means of the hourly readings from the logger were calculated for temperature, relative humidity and saturation deficit, and were denoted as *Tbar*, *RHbar* and *SDbar*, respectively. Monthly values for the Normalized Difference Vegetation Index (NDVI), were obtained from NOAA, or from Modis if NOAA figures were not available. Monthly measures were made for an area within 2km of the Research Station offices. For modelling purposes, monthly NDVI figures were used to generate daily estimates, using linear interpolation.

Data from the logger were the most useful for our purpose since, in addition to the temperature and rainfall data available from the Stevenson screen, the logger also produced measures of relative humidity and, thereby, saturation deficit. Accordingly, some of our statistical models used biological data produced between 17 November 1991 and 31 December 1999, when several meteorological variables were available from the logger, to serve as candidate explanatory variables. We could not, however, apply such models outside this period, before logger data were available. Accordingly, we also developed predictive models that used only NDVI and data available from the Stevenson screen, all of which were available during the whole study period. (**Supporting Information S1: Daily meteorological readings).**

### Capture and processing of female tsetse

All field studies were carried out at Rekomitjie Research Station (16°10′S, 29° 25′E, altitude 520 m), Zambezi Valley, Zimbabwe. Methods associated with the capture, ovarian dissection and measurement of wing length and wing fray of female tsetse have been detailed in numerous published papers (Ackley & Hargrove, 2017; English *et al*., 2016; Hargrove, 1991, 1994, 1995, 1999a, b; 2012, 2013, 2020, 2023; Hargrove & Ackley, 2015; Hargrove & Muzari, 2015; Hargrove & Van Sickle, 2023; Hargrove et al, 2018, 2019). Accordingly, the following provides only a summary of methods described elsewhere, but with fuller descriptions of the methods developed specifically for the present analyses.

### Ovarian dissection

Between September 1988 and December 1999, field-caught tsetse, collected in cages or in glass or Perspex tubes (75 x 25 mm), were placed under a moist black cloth in a polystyrene box and transferred to a field laboratory for further processing. Adult females were subjected to ovarian dissection (*ca*. 98% within 24 hours of capture) and assigned to one of eight ovarian categories, using the disposition and relative sizes of the oocytes within the ovarioles in the left and right ovaries. This procedure can be used to determine unequivocally the number of times a fly has ovulated if, and only if, she is in her first ovarian cycle – *i.e*., has ovulated fewer than four times (Challier, 1965) **(Supporting Information S2: Interpretation of ovarian dissections)**. For older flies, the number of ovulations can only be defined modulo 4 and we cannot differentiate, using only ovarian dissection data, between flies in their second and later ovarian cycle. Thus, we cannot distinguish flies that have ovulated 4 times from those that have ovulated 8, 12, 16 *etc*. times – and similar problems exist for those that have ovulated 5, 6 or 7 times.

We are specifically interested in the factors affecting the physical size of a fly, as judged by the lengths of the *in utero* egg, and/or of the wing, as measured using a binocular microscope. Egg length was measured after the egg was dissected out of the uterus. Wing length was taken as the distance between landmarks 1 and 6 as defined by Geldenhuys et al (2023). This distance is less than the total length of a wing in perfect condition. It is chosen, however, because wing tips are frequently damaged, making it impossible to obtain consistent, accurate measures of the full length.

### Temperature dependent durations of oogenesis, pregnancy and pupal development

Development rates in tsetse are strongly temperature dependent at all stages of life. Table 1 provides estimates, for female *G. m. morsitans* and *G. pallidipes*, of the temperature-dependent daily rates of passage to first ovulation, through subsequent pregnancies and during all development that occurs within the puparial case. Rates of *G. m. morsitans* pupal development were studied in detail by Phelps & Burrows (1969). No such data are available for other species of tsetse species, and we assume identical pupal development rates for *G. pallidipes*. We also assume that, as a mature oocyte is ovulated into the uterus, a new oocyte starts developing and is itself ready to be ovulated shortly after the next larviposition. For each of these developmental stages, we estimated the proportion of development completed on a given day from the mean temperature for that day, based on Table 1. The proportions completed were then accumulated over consecutive days until the first day when the estimated proportion exceeded a value of 1.0, thereby defining the estimated number of days required to complete the developmental stage in question.

**Table 1.** Estimated rates of development of different life stages of female tsetse. All rates have units of days^-1^. Temperatures (*T_bar_*) in the Phelps & Burrows (1969) study were maintained at constant levels in the laboratory. For the field estimates in this table *T_bar_* was taken as the mean of the daily maximum and minimum temperatures, measured in a Stevenson screen. The rate *r*_2_ was taken as the inverse of the observed pregnancy duration, which decreased with temperature.

|  |  |  |
| --- | --- | --- |
| Rate ($r_0$) Female | $r_0 = k_0/(1+\exp(a_0 + b_0 T_{bar}))$ | Phelps & Burrows (1969) |
| pupal development | $k_0 = 0.05689; a_0 = 5.470; b_0 = -0.2457$ | Hargrove & Vale (2020) |
| Rate ($r_1$) passage<br>to first ovulation | $r_1 = 0.1428$ | Hargrove (2012) |
| Rate ($r_2$) pregnancy<br>progression | $r_2 = a_2 + b_2*(T_{bar}-T_2)$<br>$a_2 = 0.1046; b_2 = 0.0052; T_2 = 24$ | Hargrove (1994, 1995) |

### Constructing a life history for individual female tsetse

We aimed to estimate meteorological conditions prevailing when individual flies were at various well-defined stages of development, particularly during oogenesis and pregnancy when these conditions should maximally impact the size of the developing immature fly, via effects on their mothers. This aim is achieved, using ovarian dissection and meteorological data, as described below. If we know how many times a fly has ovulated, we can estimate its age to the nearest 9-12 days, or to the nearest 3-4 days if we also use the lengths of any larva *in utero* and of the two largest oocytes in the ovaries. This information allows computation of the proportion of pregnancy completed and, given knowledge of the inter-larval period, the estimated number of days since the last ovulation (Hargrove, 1994, 1995). For a fly of known chronological age, and date of capture and dissection, we used temperature profiles over the period of the fly’s life, to reconstruct its life history – using the temperature-dependent rates of development (Table 1) during various phases of the tsetse life cycle. This procedure is best illustrated using a specific example.

### Winding back the clock

Consider a female *G. pallidipes*, captured on 17 July 1993 and found on examination to be in ovarian category *c* = 3. The lengths of the fly’s uterine and ovarian contents were used to estimate that 0.58 of the current pregnancy had been completed – *i.e*., the fly had completed 3.58 ovulations. On 17 July 1993 the mean temperature was 20.9°C and the equation for *r*_2_ in Table 1 predicts that 0.089 of an ovulation cycle was completed on that day, so the fly was estimated to have completed 3.580 - 0.089 = 3.491 ovulations at the start of the previous day, 16 July 1993. The *r*_2_ rate equation, and daily mean temperatures, were used to estimate the proportion of pregnancy completed on each preceding day until the predicted ovulations completed first takes a value ≤ 1.0, which was then taken as the day after the female ovulated for the first time. This event occurred between 6 June, taken as the first day of *c* = 1, and 5 June, taken as the day at the end of which the female ovulated for the first time. The rate *r*_1_ in Table 1 was then used in like manner to estimate that the first day of adult life occurred on 30 May. The adult female thus emerged from its puparial case on 29 May (Table 2) – at the end of which day the pupa was taken to have completed its pupal life. To find the day on which pupal life started, *i.e*., the date of larviposition, we continued the rewinding procedure, using now the temperature-dependent rate *r*_0_ of female pupal development (Table 1). This resulted in a prediction that pupal development started on 26 April, so that 25 April marked the last day of pregnancy, with larviposition estimated to have occurred at the end of that day (Table 2). Finally, we repeated the rewinding procedure, again using the rate *r*_2_, to estimate that the pregnancy producing our example fly spanned the dates 16 – 25 April, and that the period of oogenesis was 7 – 15 April (Table 2). (**Supporting Information S3. *Example calculation.xlsx*** for all details of above calculations).

**Table 2.**
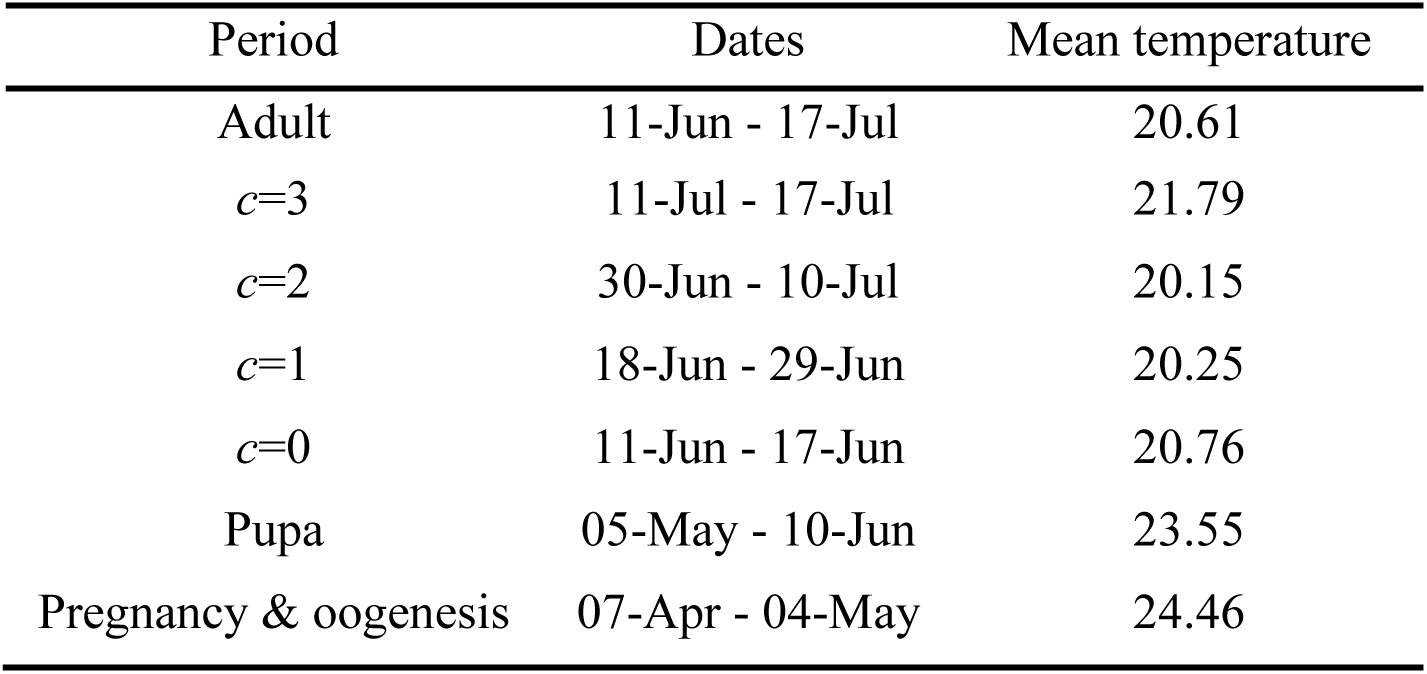
Mean daily temperatures during different stages of the development of a female *G. pallidipes* captured on 17 July 1993, when she had ovulated three times and had completed an estimated 58% of the current pregnancy. The rows labelled *c*=0, 1, 2, 3 denote the periods spent between successive ovulations.

| Period | Dates | Mean temperature |
| --- | --- | --- |
| Adult | 11-Jun - 17-Jul | 20.61 |
| $c=3$ | 11-Jul - 17-Jul | 21.79 |
| $c=2$ | 30-Jun - 10-Jul | 20.15 |
| $c=1$ | 18-Jun - 29-Jun | 20.25 |
| $c=0$ | 11-Jun - 17-Jun | 20.76 |
| Pupa | 05-May - 10-Jun | 23.55 |
| Pregnancy & oogenesis | 07-Apr - 04-May | 24.46 |

The same procedure was carried out for all females in their first ovarian cycle – *i.e*., those that had ovulated 0, 1, 2 or 3 times. For flies in their second and later ovarian cycles, ovarian dissection alone cannot be used to say whether a fly is in its second, third or later ovarian cycle. Such flies can only be placed in *c* = 4+4*n*, 5+4*n*, 6+4*n* or 7+4*n* where *n* can, in principle, take any non-negative integer value (Challier, 1965). Wing fray can, in principle, be used to distribute these older flies between the second, and later, ovarian cycles flies. Applying the procedure to all such flies clearly leads, however, to errors of allocation (Hargrove, 2020). Accordingly, we only produced full life histories for flies in ovarian categories *c* = 0 to 3 – where the problem of age uncertainty does not arise. (**Supporting Information S4. Life histories for qualifying adult female *G. pallidipes*.xlsx)**

### Wing and egg lengths

#### Temporal pooling of wing lengths

The procedure described in the previous section identifies daily cohorts of female tsetse, which were deposited as pupae on the same day. Figure 1 illustrates this procedure for flies of different ages, denoted by red arrows of different lengths, all of which emerged on day 1 of a week. By assumption, these flies experienced identical meteorological conditions throughout their development – *i.e*., during oogenesis, their mother’s pregnancy and as pupae. Ideally, one could now relate the mean wing sizes of daily cohorts to meteorological and other environmental variables. Time series of daily mean wing size were, however, quite noisy due to the large size variability among individual flies, and to small, sometimes zero, catches related to individual days of larviposition. Accordingly, we also pooled flies, based on their larviposition dates, into both monthly and weekly groups. We used monthly pooling to graphically explore coarse annual cycles of fly size and environmental change. Weekly pooling captured rapid changes occurring during some parts of annual cycles. We also used weekly averages from these groups, in our statistical modelling of *G. pallidipes* size versus environmental variables. For *G. m. morsitans* modelling, we used averages from monthly groups, due to our much smaller sample size for this species.

**Figure 1.**
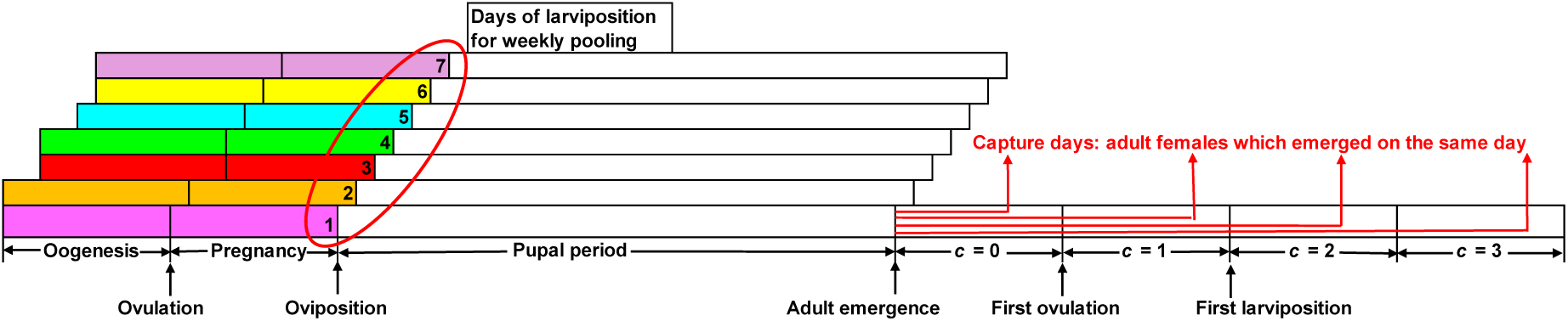
Diagrammatic representation of the procedure for pooling female tsetse, based on their estimated life histories. Examples are given for four sampled flies, found on dissection to be in ovarian categories *c* = 0, 1, 2 and 3, respectively – all of which are estimated to have emerged as adults on the same day. All flies within the red oval can be pooled into one weekly group, based on the end dates of their wing-size developmental period: analogous procedures generated data pooled by month.

Our weekly pooling approach, illustrated in Figure 1, is based on the life histories of all flies that larviposited over a 7-day period (Monday through the following Sunday) denoted by the numbers in the red oval in Figure 1. All larvae deposited on a given day are assumed to experience identical meteorological conditions during the pupal period and to emerge as adults on the same day. The lengths of the red arrows indicate the ages at capture of all flies estimated to have been larviposited on day 1 of the week. An analogous situation applies to each day of the week. Weekly means of wing length were calculated as unweighted means, across all flies larviposited during the week.

### Environmental averaging

Our wing size analyses assume that a fly’s eventual size is entirely determined by environmental conditions during the oogenesis and maternal pregnancy stages (Fig. 1). Accordingly, for each fly, we calculated the unweighted mean of daily values, separately for each available meteorological variable, over that fly’s total time interval for those two stages (Table 2, Fig. 1). These averaged variables were then used in exploratory correlations and regressions between individual fly sizes and their associated environmental regimes. For weekly and monthly groups of flies, we further averaged the per-fly means of environmental variables, across all flies within each group. This provided weekly and monthly average values of environmental variables, which we then related to corresponding group averages of fly size.

### Egg lengths

We assume that the length of an egg in a female’s uterus depends only on conditions she experienced during the oogenesis period resulting in the production of that egg – *i.e*., the ovulation cycle most recently completed prior to the capture date. We do not, therefore, need a fly’s full life history to determine the oogenesis period of the egg it carries *in utero*. To see this refer to Figure 1, and the three example flies captured where *c* = 1, 2 and 3, respectively. For the fly captured in *c* = 3, we only need to generate the start and end dates for *c* = 2 and analogous statements apply to the flies captured in *c* = 2 and *c* = 1. We could thus accurately determine development periods for eggs carried by females *c* = 4+4*n* to 7+4*n*, even without a reliable estimate of those females’ chronological ages. Accordingly, we included eggs from females of all ovarian categories in our analyses of egg lengths.

We also pooled eggs on a weekly or monthly basis, in similar fashion to the pooling done for wing size analyses. Egg-bearing females, and their eggs, were pooled into groups whose end dates of their eggs’ oogenesis stage all fell within the same week (or month). Environmental variables were averaged over the oogenesis period for each egg, and these averages were again averaged over each pooled subset of eggs, in the same fashion as for wing size analyses. Maternal wing length was also averaged across pooled subsets, as another candidate explanatory variable.

### Sample sizes

We used subsets of data from a study carried out between September 1988 and December 1999, during which period 151,269 female *G. pallidipes* and 19,569 female *G. m. morsitans* were subjected to ovarian dissection and provided both a valid ovarian category and a wing length (Table 3, row 1). Throughout the study, daily maximum and minimum temperatures, and rainfall were available from the Stevenson screen and monthly measures of NDVI were accessed from the internet. Relative humidity and saturation deficit were available only from 17 November 1991, for 64% of the dissected *G. pallidipes* and 49% of the *G. m. morsitans* (Row 3). About 51% of all *G. pallidipes*, and 54% of all *G. m. morsitans*, had ovulated fewer than four times and could therefore provide full life histories. We estimated chronological age of each such fly and the period it had spent within its mother during oogenesis and pregnancy (Row 4). Among these flies, about half of each species spent the developmental period at times when meteorological data, including relative humidity, were available from the logger (Row 6).

**Table 3.** Numbers of adult female tsetse available for analysis of factors affecting wing and egg lengths in *G. pallidipes* and *G. m. morsitans* sampled at Rekomitjie. Flies had a full life history only if they were in ovarian categories *c* = 0 to 3 and if the wing length was recorded. SS indicates that meteorological data were available only from the Stevenson screen; SS/L that they were also available from the logger. *n*/*d* indicates the rows used for numerator and denominator, respectively.

| Row | Wing length | n/d | *G. pallidipes* | *G. m. morsitans* |
| --- | --- | --- | --- | --- |
| 1 | Total sample |  | 151,269 | 19,569 |
| 2 | 13-Sep-88 - 16-Nov-91 SS | 2/1 | 53,954 (36%) | 10,083 (52%) |
| 3 | 17-Nov-91 -17-Dec-99 SS/L | 3/1 | 97,315 (64%) | 9486 (48%) |
| 4 | Full life history available ($c < 4$) | 4/1 | 76,871 (51%) | 10,616 (54%) |
| 5 | 13-Sep-88 - 16-Nov-91 SS | 5/4 | 28,596 (37%) | 6100 (57%) |
| 6 | 17-Nov-91 -17-Dec-99 SS/L | 6/4 | 48,275 (63%) | 4516 (43%) |
| **Egg length** |  |  |  |  |
| 7 | Total sample | 7/1 | 66,150 (44%) | 6079 (32%) |
| 8 | 13-Sep-88 - 16-Nov-91 SS | 8/7 | 24,943 (38%) | 3318 (55%) |
| 9 | 17-Nov-91 -17-Dec-99 SS/L | 9/7 | 41,207 (62%) | 2761 (45%) |

### Statistical methods

#### Relating wing size to environmental factors

We used linear correlation and multiple linear regression to model wing and egg sizes, using averaged environmental variables as potential explanatory variables. We developed our final predictive models for pooled groups, by using the Akaike Information Criterion (AIC) to help select best-fitting, most parsimonious models (Venables & Ripley, 1997).

## Results

### Meteorological patterns at Rekomitjie during the study period

The general pattern of meteorological changes at Rekomitjie is shown in Figure 2A – and in detail for July 1993-June 1994 in **Supporting Information S5**, Fig. S5.1. Temperature and saturation deficit increased steadily, and relative humidity decreased, from July until the rains started in late November/December. Temperatures then fell by 5-10°C but remained at fairly high levels until the end of the rains in March. Temperature and humidity then declined until July, with saturation deficit increasing proportionately. NDVI, which reflects photosynthetic activity and thus vegetation cover, showed roughly the same seasonal pattern as relative humidity and was invariably at a minimum at the end of the hoy dry season.

**Figure 2.**
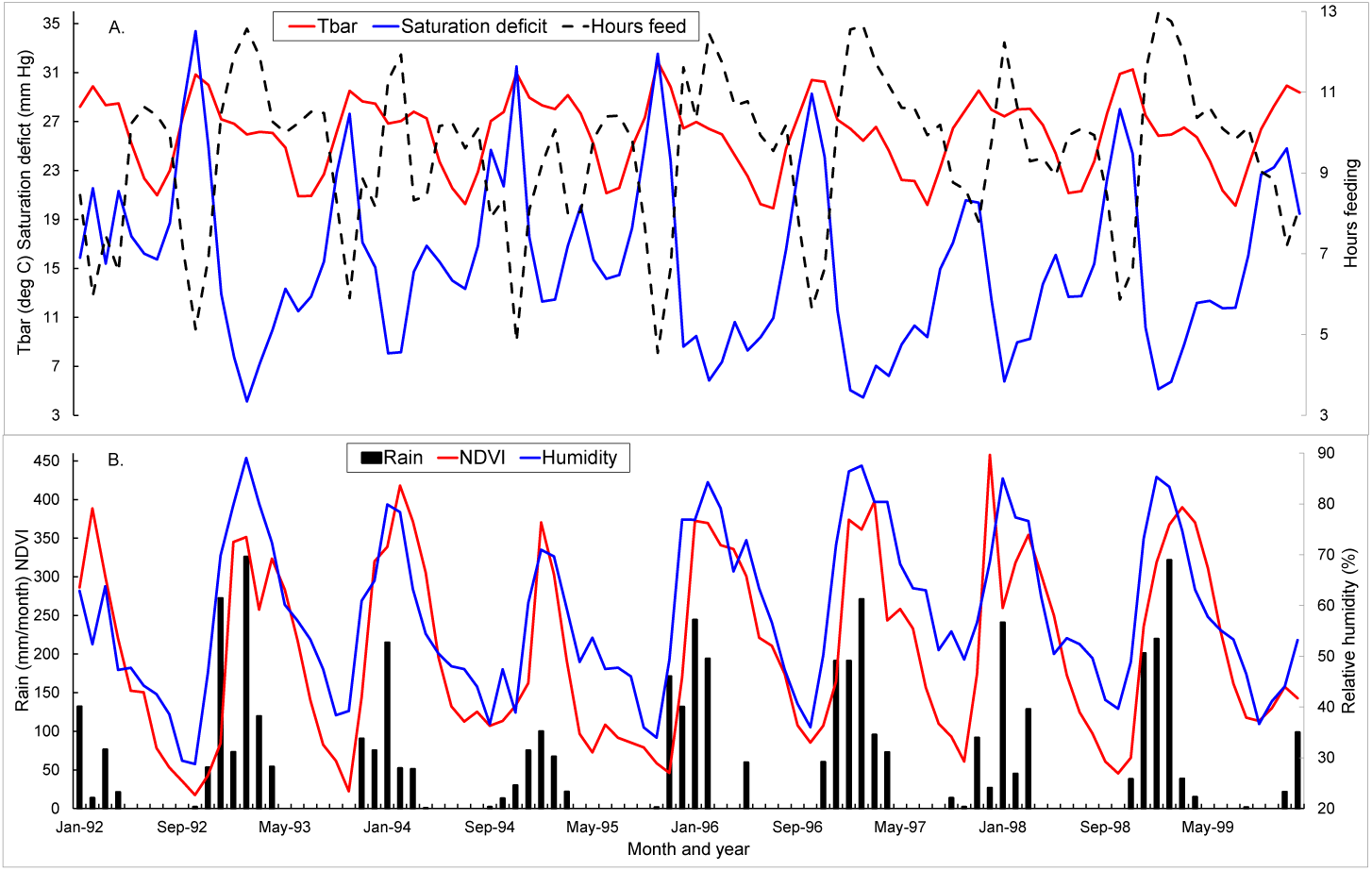
Seasonal changes in meteorological profile 1992-1999. A. Monthly variation in daily mean temperature (*Tbar*; °C), saturation deficit (mB) and the estimated number of hours per day suitable for tsetse feeding activity. B. Total rainfall per month and mean monthly values for the Normalised Difference Vegetation Index (NDVI) and relative humidity over the same period. Values for NDVI, which varied between 0.175 and 0.804, have been scaled for purposes of display.

When days were viewed singly, the numbers of hours suitable for feeding varied between 1 and 13 hours (**Supporting Information S6**, Fig. S6.1). When averaged by month the value varied between about 7 and 13 (Fig. 2A). Mean hours suitable for feeding were longest in January and February (11 hours) and shortest in September – November (8, 6.5 and 7 hours, respectively). For all other months the mean was 9-10 hours (**Supporting Information S6**, Fig. S6.2). Months with the lowest number of hours suitable for feeding were also those in which temperature and saturation deficit were highest, and relative humidity and NDVI were lowest (Fig. 2).

### Seasonal variation in the durations of oogenesis, pregnancy and pupal period

Our life history estimates generated the durations of pregnancy and pupal life for all young flies (*c* < 4) starting either of these processes on any day between 17 November 1991 and the end of 1999. All durations were shortest in November and longest in July of each year (Fig. 3). Over the period of analysis, pregnancy duration for *G. m. morsitans* and *G. pallidipes* varied from 7 to 14 days (Fig. 3A), but 84% of oogenesis and pregnancy durations were in the interval 8 to 11 days, inclusive (Fig. 4A, B). Pupal durations, likewise, varied over the large range of 19-57 days, but 70% lay in the interval 23-33 days inclusive (Fig. 4C).

**Figure 3.**
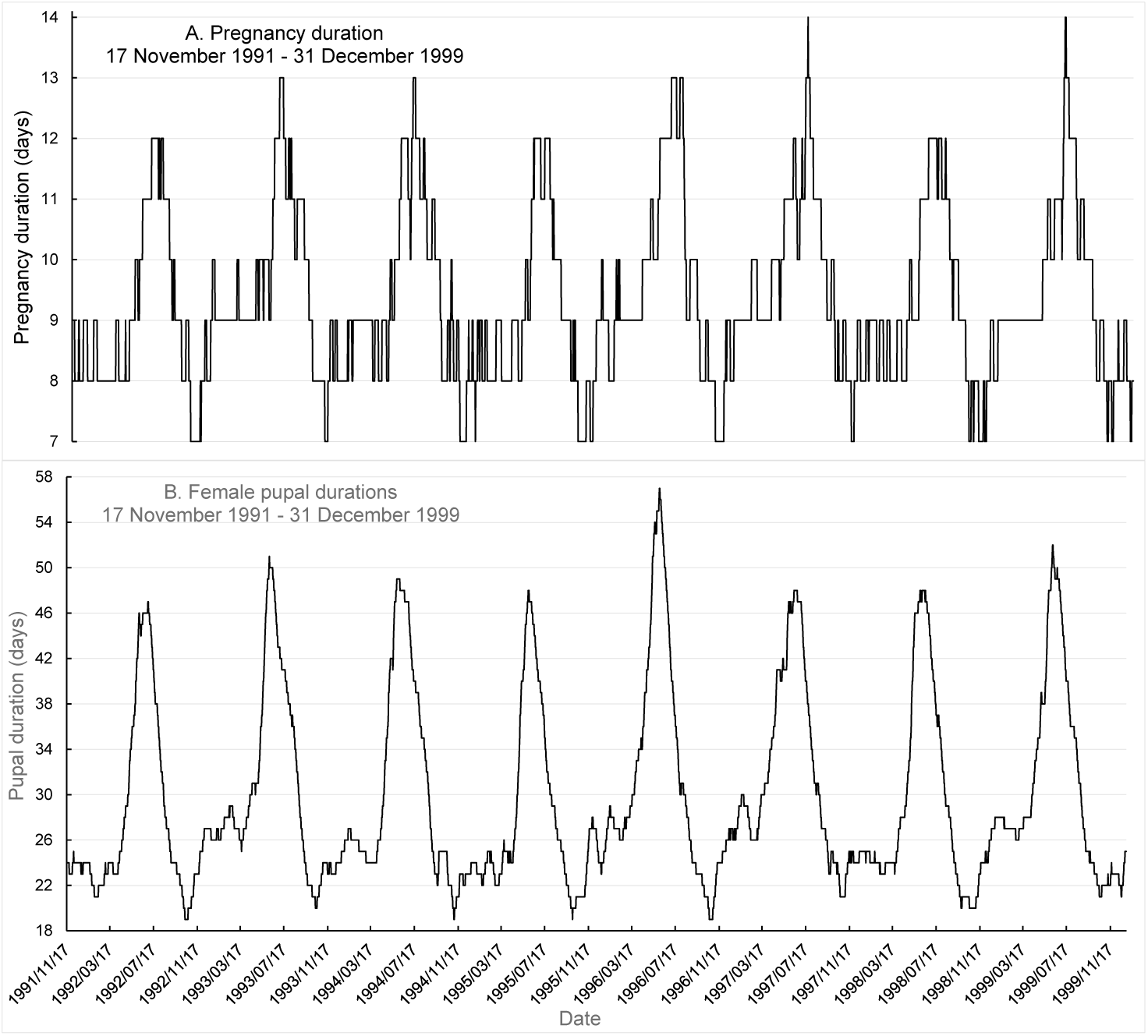
Estimated durations of: A. Pregnancy; B. Female pupal duration, for *G. m. morsitans* and *G. pallidipes* by day of larviposition in typical larviposition sites, at Rekomitjie, November 1991 – December 1999.

**Figure 4.**
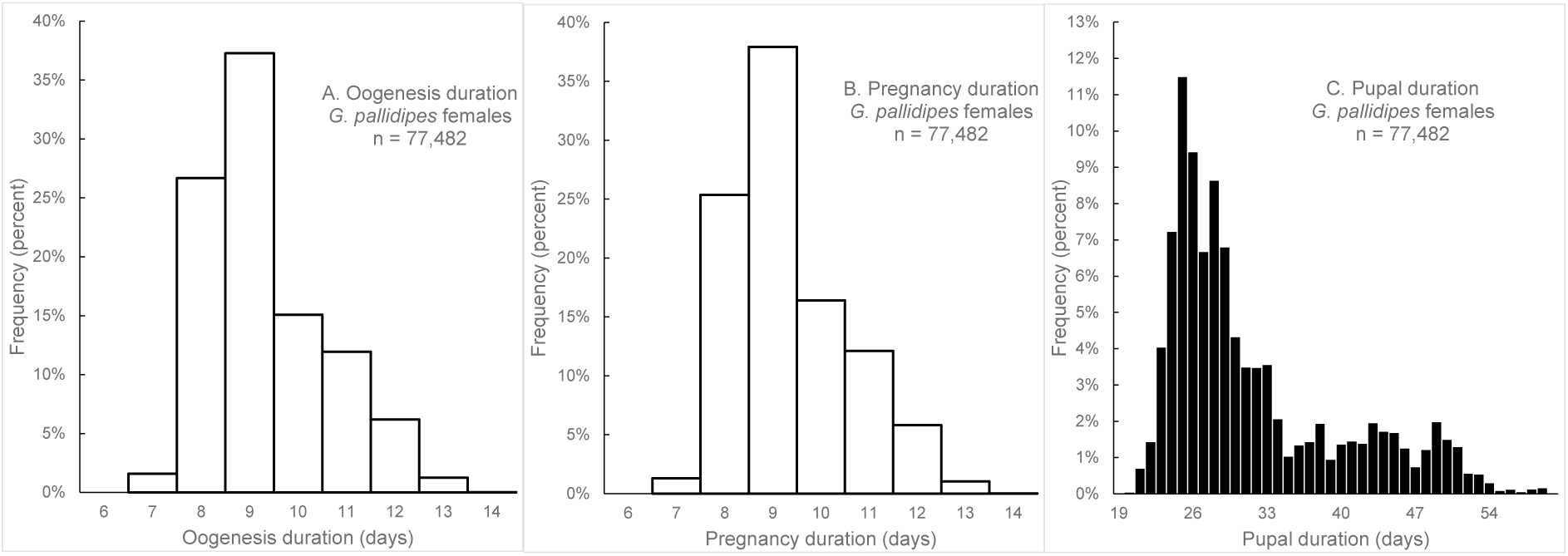
Durations of (A) oogenesis (the time that the largest oocyte spends in the ovary), (B) pregnancy and (C) pupal life – all calculated from the estimated times of the starts and ends of oogenesis, pregnancy and pupal development of individual flies in ovarian categories *c* = 0-3.

The age distribution of all *G. pallidipes* females for which we had a life history shows an early peak of activity immediately following adult emergence (Fig. 5) followed by a hiatus in catches until the age of 7-8 days – when females ovulate for the first time. Lower frequencies at ages 2-6 days may be attributed to reduced activity in flies that are using their first two blood meals to build thoracic flight muscle (Bursell, 1961; Hargrove, 1975, 1999a, b; Vale *et al*., 1976). Frequencies change little for ages 8-26 days, then fall away because significant proportions of flies have ovulated for a fourth time: we could not confidently estimate their chronological age and did not, therefore, generate their life histories.

**Figure 5.**
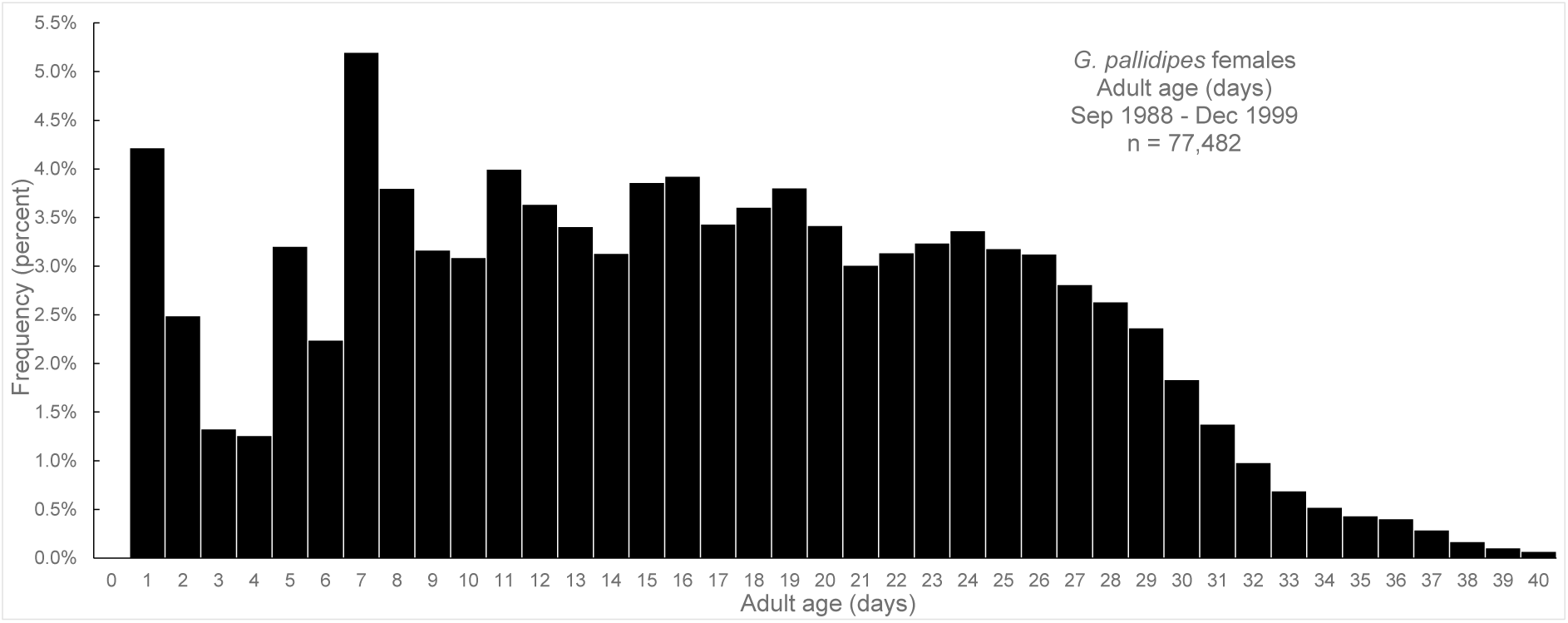
Distribution of ages of female *G. pallidipes*, estimated by ovarian dissection. Note that ages are only estimated for flies in *c* = 0-3.

### Distributions of egg and wing lengths

Egg and wing lengths are both greater in *G. pallidipes* than *G. m. morsitans* but the difference is much less marked in eggs (**Supporting Information File S7, Figures S7.1, 7.2).** Samples were excluded from analyses if it seemed likely that the fly had been identified as the incorrect species, or if there had clearly been an error in the measure of the egg or wing length. This happened in <1% of wings measured, and <0.1% for eggs (**Supporting Information** Table S7.1).

### Seasonal changes in egg and wing lengths

Mean egg and wing lengths of female *G. pallidipes* and *G. m. morsitans* pooled by month of capture, varied in a highly regular fashion with season – with a minimum in December at the end of the hot dry season and beginning of the rains (Fig. 6). Thereafter, mean wing length increased rapidly, peaking in about April/May in the warm dry season, then declined again rapidly between August and December. There was a similar regularity in the changes with time in mean wing lengths of flies estimated to have emerged in each month – but the cycle of changes lagged by about a month behind the length changes for flies captured in that month. The same pattern of length changes was evident every year (**Supporting Information File S8. Figure S8.1)**.

**Figure 6.**
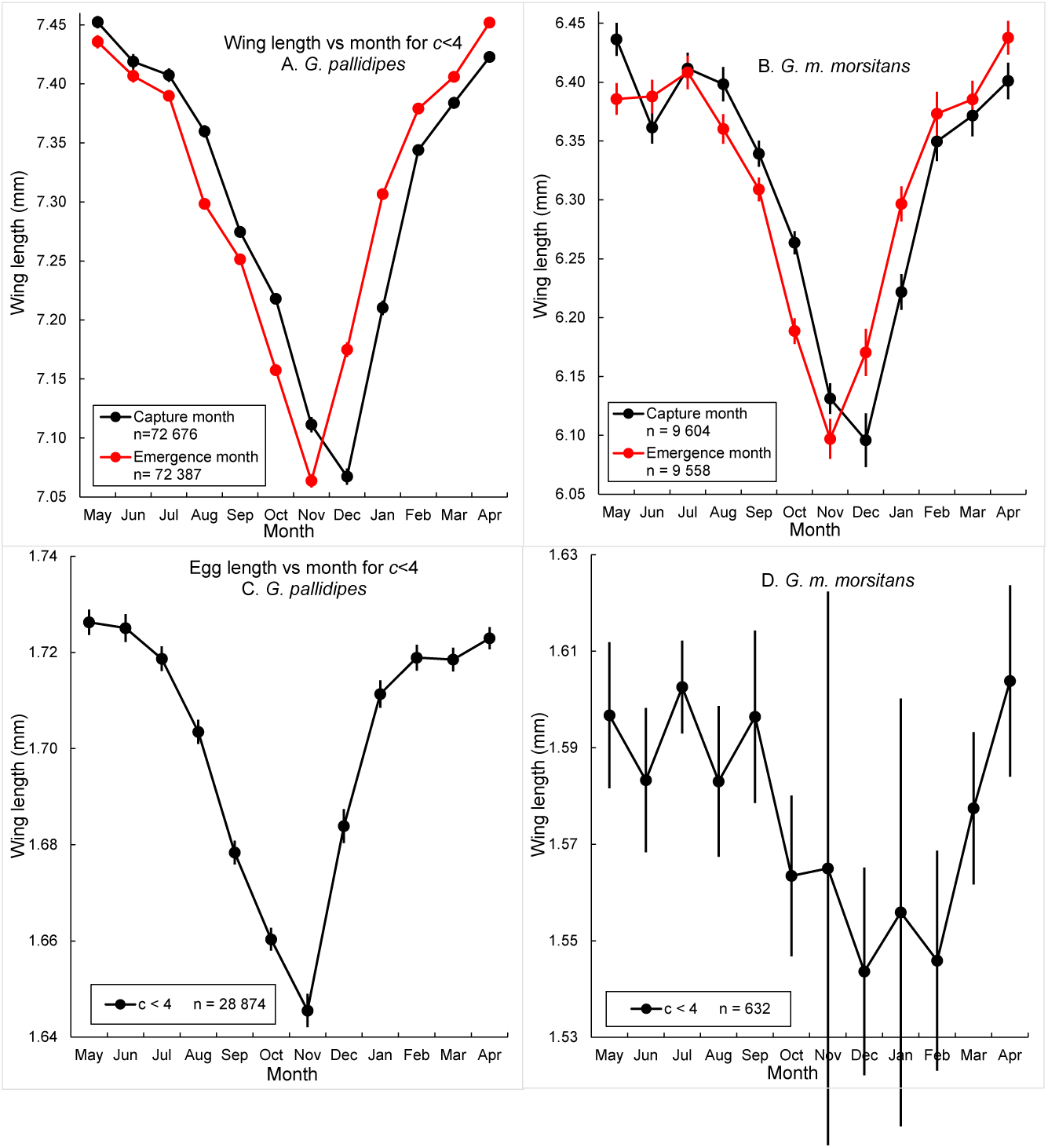
Mean wing and egg lengths of *G. pallidipes* (A, C) and *G. m. morsitans* (B, D) sampled at Rekomitjie between Sep 1998 and Dec 1999. Lengths are plotted as a function of the month of capture of the adult female, and wing lengths also against the month in which females were estimated to have emerged as adults. Catches are pooled over all years and restricted to females that had ovulated fewer than four times.

We pooled wing and egg lengths on year of capture, then estimated the mean lengths as a function of month and of ovarian age category (Fig. 7). Mean wing lengths for both *G. pallidipes* and *G. m. morsitans*, of all ovarian categories, declined sharply between June and December, most particularly for flies where *c* = 0 (Fig. 7 A, B). For egg lengths, the picture was even clearer, with the eggs of youngest mothers (*c* = 1) being significantly shorter, in every month of the year, than eggs of all older flies (Fig. 7 C, D). The eggs of *G. pallidipes* female where *c* = 2 were also significantly shorter than for older flies, throughout the year. The difference was less marked for *G. m. morsitans*, reflecting in part the smaller sample sizes for this species. Age effects are seen more clearly in **Supporting Information File S8, Fig S8.2**, where egg lengths are plotted against ovarian category for data stratified by month of maternal capture.

**Figure 7.**
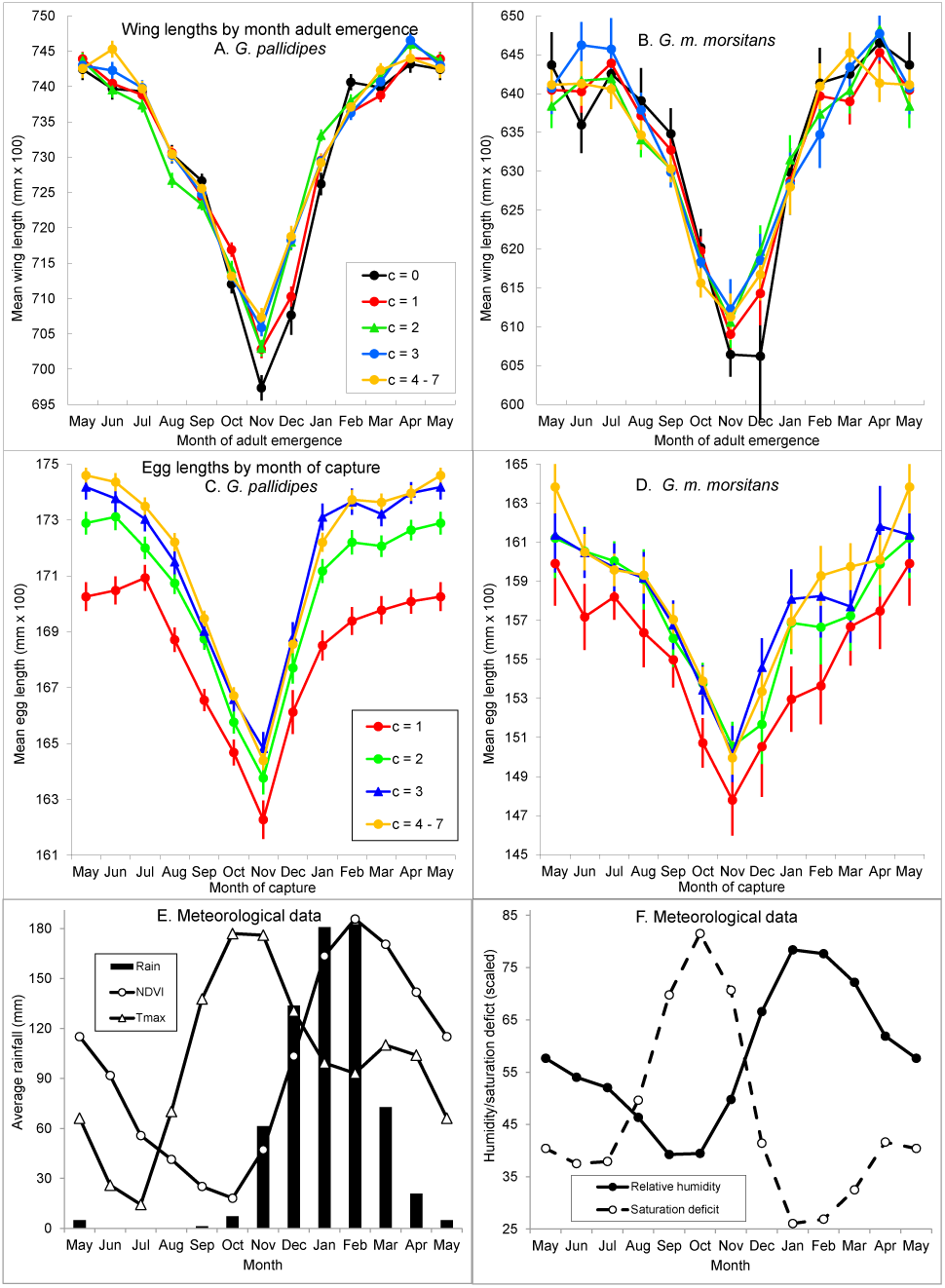
Mean wing and egg lengths for females of *G. pallidipes* (A, C) and *G. m. morsitans* (B, D). Eggs were pooled by the month of their mother’s capture; adults on the estimated month of their emergence, which could only be estimated for flies where *c*<4. (E) Mean monthly values of mean daily temperature (*T_bar_*), NDVI and rainfall. (D). Relative humidity and saturation deficit were available only from November 1991. NDVI and *T_bar_* in (E) and saturation deficit in (F) were scaled to show relative timing of changes with season of all five variables. Minimum and maximum monthly values for the scaled variables were: NDVI (0.225, 0.664); *T_bar_* (20.88, 29.84); saturation deficit (8.68, 27.17).

### Wing length as a function of age and of month of either adult emergence or of capture

#### Month of emergence as an adult

Wing lengths for females of both species that emerged within the week prior to capture (*c*=0), declined steadily from July – November, then increased rapidly until February, with smaller change between February and July (Fig. 6 A, B). Within most months of the year, mean wing length varied little with ovarian category *c*, consistent with the implicit assumption that the wing length of an individual adult tsetse does not vary with age. In November and December, however, mean length increased markedly with *c*. These changes can be seen more clearly when wing lengths of female *G. pallidipes* were plotted against *c*, for each month of the year in which the fly was estimated to have emerged as a teneral adult (**Supporting Information File S8,** Fig S8.3).

The increases in wing length with age, during the hot season, are consistent with disproportionately high losses among small, young, flies in hot-dry months, so that mean sizes of surviving older flies were thereby increased. This accords with published evidence indicating selection against small individuals in several different tsetse species (Jackson, 1948b; Bursell & Glasgow, 1960; Phelps & Clarke, 1974; Dransfield *et al*., 1989). Previous authors all concluded, however, that losses of small flies were restricted to the teneral period, *i.e*., only in very young flies. By contrast, Figure S7.3 suggests that the size of flies emerging as adults in October through December continues to increase for some weeks after flies would have taken their first blood meal. We investigated the discrepancy further, using flies whose ages were estimated to the nearest day. In Figure 8A – C, note that from July through September wing lengths were getting shorter each month, but there was little change in wing length with age, within each month. In October and November, the wing lengths of the youngest flies continued to get progressively smaller (Fig. 8A, B); but there was then a more sustained increase in wing length with age within each month, with the November increase continuing for more than three weeks. In December (Fig. 8C), when the rains arrived and temperatures dropped, wing length started from a higher base in young flies, but there was still a sustained increase with age over the first three weeks of life. This trend was weakly sustained in January (Fig. 8D); in February (Fig. 8E), changes with age were restricted to flies in the first week of life. The same was true for April and May (Fig. 8D, E); for March and June there was little change with age (Fig. 8F).

**Figure 8.**
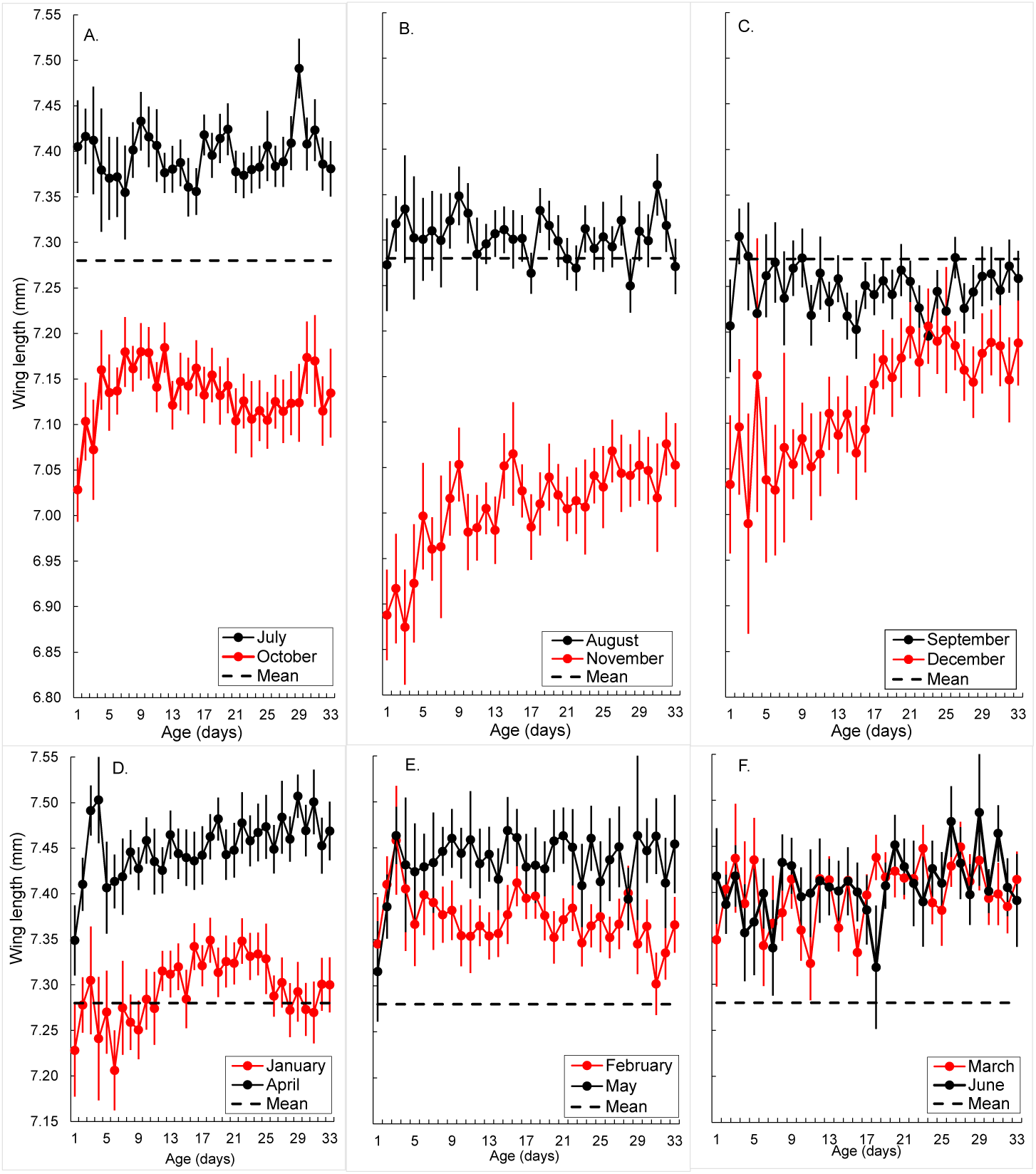
Wing length in female *G. pallidipes* sampled at Rekomitjie between Aug 1988- Dec 1999, as a function of the flies’ daily ages and months in which the flies were estimated to have emerged as teneral adults. Horizontal dashed lines indicate the overall mean wing length, across all flies. Total sample size; 76,506.

**Figure 8.**
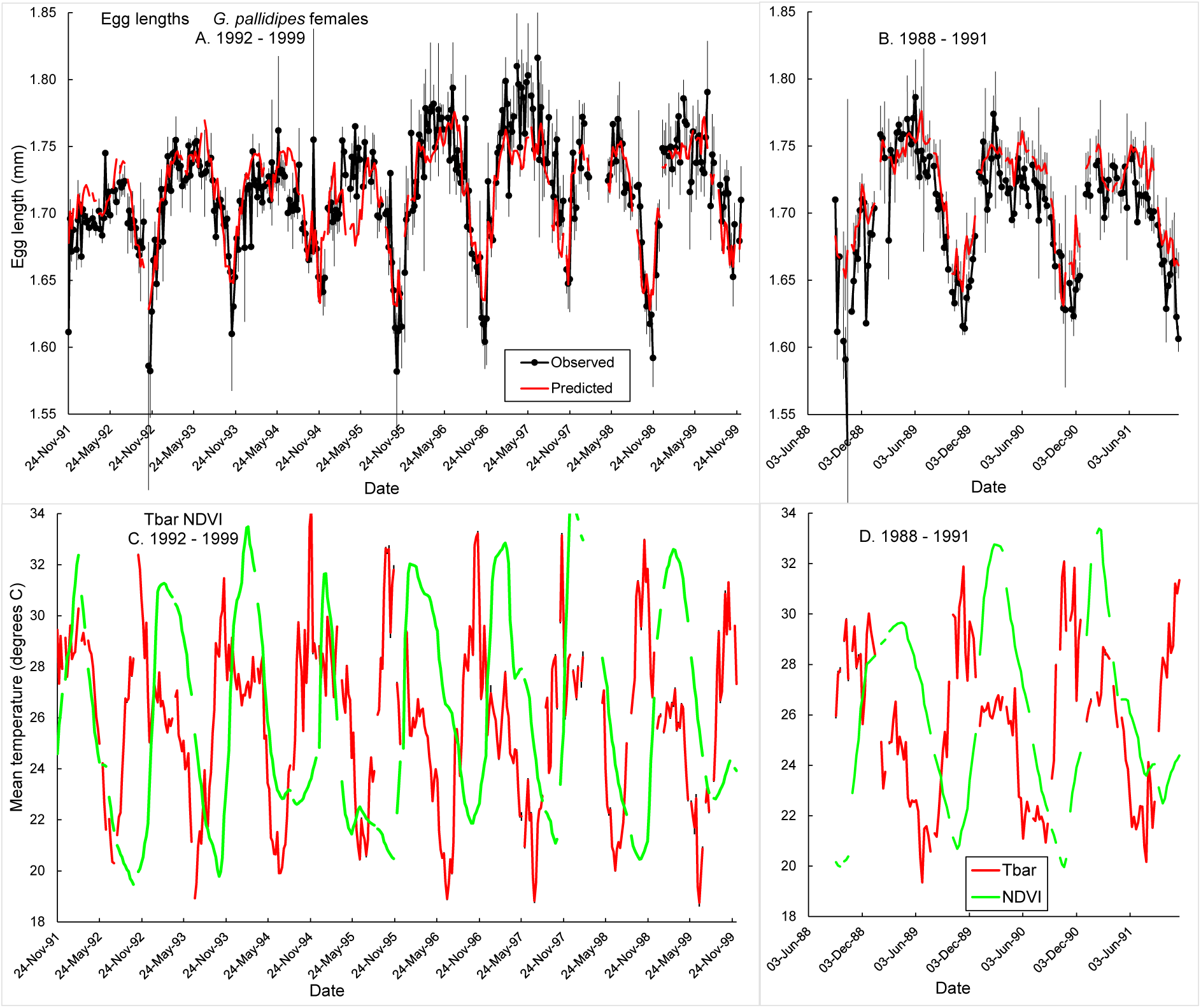
Egg lengths of *G. pallidipes* sampled at Rekomitjie. Observed weekly mean egg lengths, with 95% confidence intervals, for flies pooled weekly based on the end of oogenesis, for A. December 1991 – December 1999: B. November 1988 – November 1991. Predicted lengths in A result from fitting the model shown in Table 5A. Predicted weekly mean lengths in B are true predictions from the model that was fitted to the data in A. C, D. Weekly mean temperatures and NDVI calculated over the period of oogenesis leading to the production of eggs in A and B, respectively.

**Table 5.** Regression of egg length (*ulmm*) in tsetse as a function of mean temperature (*Tbar.*wkmn) and NDVI (*ndvi.*wkmn).

| Coefficients | Estimate | Std. Error | t value | Pr(>\|t\|) |
| --- | --- | --- | --- | --- |
| (Intercept) | 1.871 | 0.0110 | 170.5 | <2e-16 *** |
| *Tbar.wkmn* | -0.00831 | 0.000397 | -20.9 | <2e-16 *** |
| *ndvi.wkmn* | 0.136 | 0.00792 | 17.2 | <2e-16 *** |

| Coefficients: | Estimate | Std. Error | t value | Pr(>\|t\|) |
| --- | --- | --- | --- | --- |
| (Intercept) | 1.825 | 0.0383 | 47.7 | < 2e-16 *** |
| *Tbar.momn* | -0.0105 | 0.00132 | -7.97 | 4.41e-11 *** |
| *ndvi.momn* | 0.0471 | 0.0234 | 2.02 | 0.048 * |

Residual standard error: 0.0306 on 62 degrees of freedom
Multiple R<sup>2</sup>: 0.542, Adjusted R<sup>2</sup>: 0.527; *F*-statistic: 36.7 on 2 and 62 DF, *p*-value: 3.056e-11

#### Month of capture

When flies were grouped with respect to their month of capture, rather than emergence, there were more striking age-related changes in wing length. From August to December, mean length increased sharply with ovarian age (Fig. 9A), whereas in January through April, the mean length decreased with age (Fig. 9B). These changes are readily understood as a natural consequence of the seasonal changes in the wing lengths of emerging adults. For example, the oldest flies among those sampled in December would have been deposited as larvae in the cool dry season, when the flies are biggest, while the youngest flies would have been deposited at the height of the hot dry season when they are smallest. Likewise, in February, the youngest flies will have been deposited in January and February, when relative humidity is highest and temperatures are declining (Fig. 2) – whereas flies of ovarian ages 4+4*n* – 7+4*n* will have been deposited in September-November, when temperatures are highest and relative humidity lowest. Similar analyses carried out on the data available for *G. m. morsitans* provided a generally similar picture – based on much smaller samples size with, accordingly, greater variability (**Supporting Information S9**, Figure S9.1, C, D). The steady decrease in wing and egg lengths in the second half of each year coincides with the onset of the hot dry season in the Zambezi Valley and increases in length began with the onset of the rains (Fig. 7E, F). These results raise the question of which meteorological features are best correlated with the observed changes in egg and wing length.

**Figure 9.**
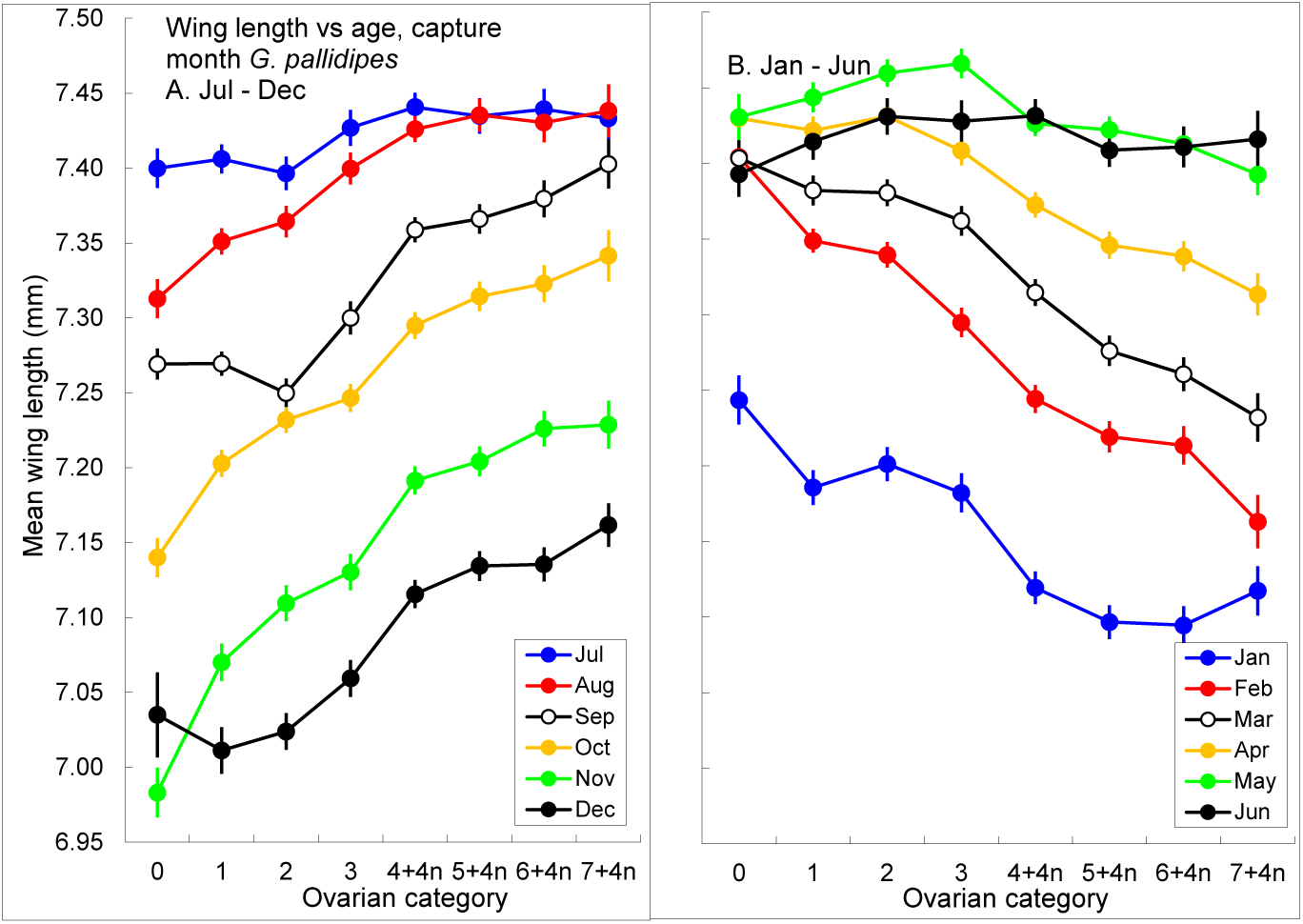
Wing length in female *G. pallidipes* sampled at Rekomitjie between Aug 1988 and December 1999, as a function of the ovarian age of the fly and the month in which the fly was captured. Total sample size; 151,789.

**Figure 9.**
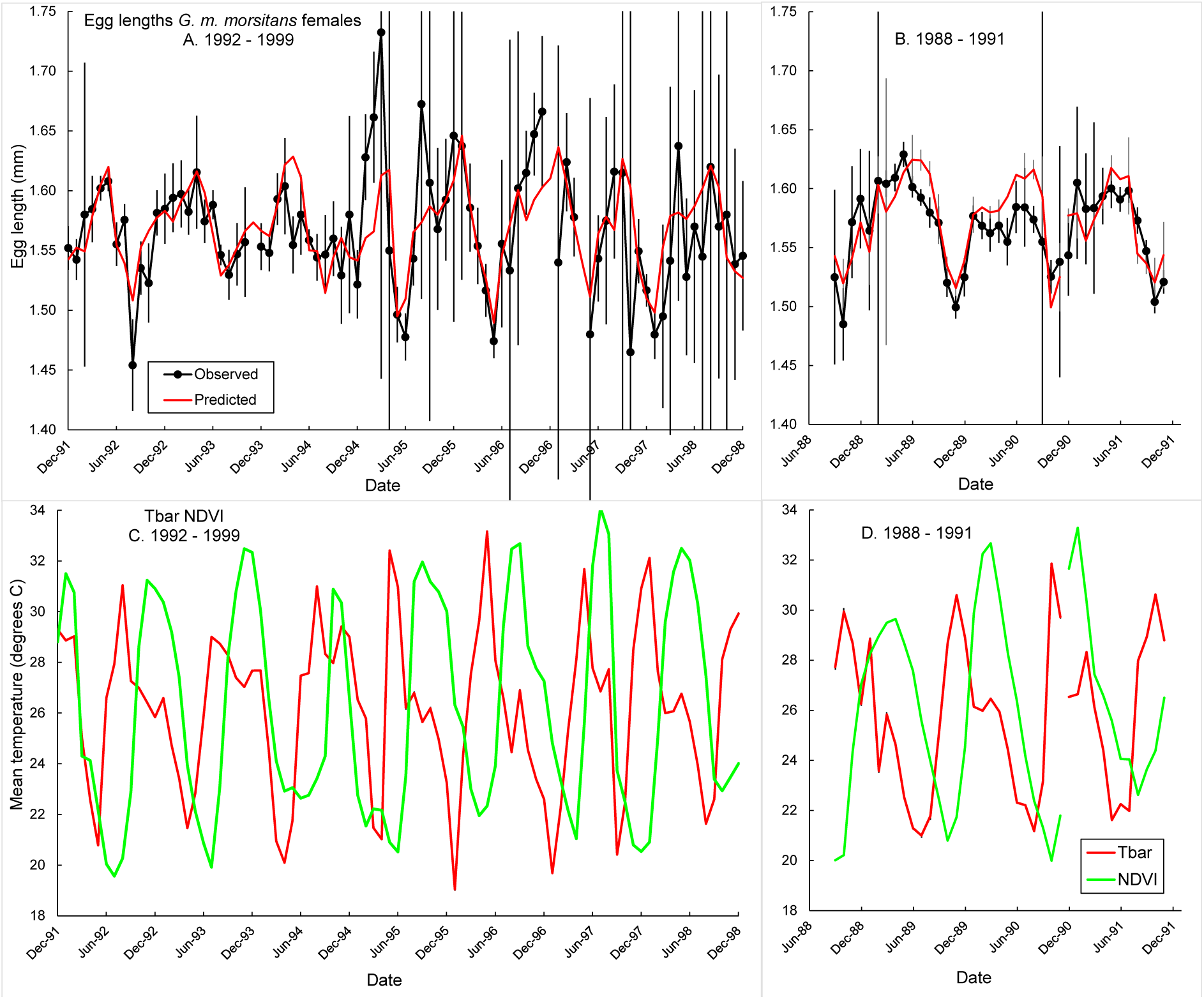
Egg lengths of *G. m. morsitans* sampled at Rekomitjie. Observed monthly mean egg lengths, with 95% confidence intervals, pooled monthly based on the end of oogenesis, for A. December 1991 – December 1999: B. November 1988 – November 1991. Predicted lengths in A result from fitting the model shown in Table 5B. Predicted monthly mean lengths in B are true predictions from the model that was fitted to the data in A. C, D. Temperatures and NDVI calculated over the period of oogenesis leading to the production of eggs in A and B, respectively.

**Figure 9.**
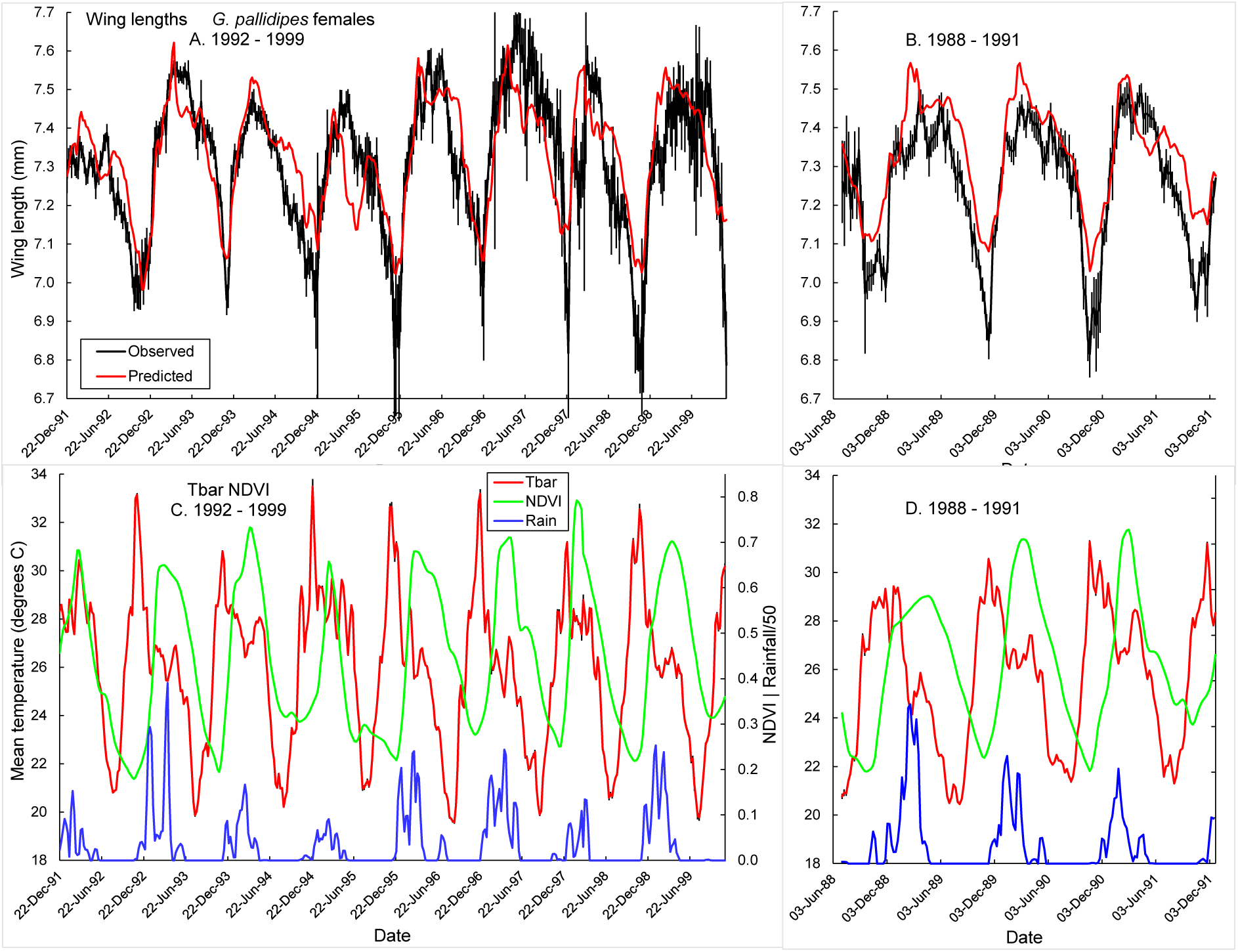
Observed weekly mean wing lengths of *G. pallidipes* females at capture (black lines, with 95% confidence intervals) plotted against the week of their larviposition. Red lines show lengths predicted by the model detailed in Table 6. (A) Flies captured 17 Nov 1991 - 31 Dec 1999; (B) 13 Sep 1988 - 16 Nov 1988. Red, green and blue lines in Fig 10C and D show corresponding weekly mean temperatures, NDVI and rainfall, respectively, averaged over the periods of oogenesis and pregnancy – *i.e*., while the fly was developing within its mother’s ovaries and uterus, respectively.

**Table 6.** Regression of mean weekly wing length (*wlmm*) in *G. pallidipes* as a function of the mean temperature (*Tbar.*wkmn), NDVI (*ndvi.*wkmn) and rain (*rain.*wkmn). Wing lengths are the mean values for flies caught in consecutive one-week (7-day) periods; temperatures, NDVI and rain are the estimated mean values of these variables over the period when the captured flies were developing in the ovaries and uteri of their mothers.

| Coefficients: | Estimate | Std. Error | t value | Pr(>\|t\|) |
| --- | --- | --- | --- | --- |
| (Intercept) | 7.564 | 0.0482 | 157.0 | <2e-16 *** |
| *Tbar.wkmn* | -0.02267 | 0.00173 | -13.1 | < 2e-16 *** |
| *ndvi.wkmn* | 0.7314 | 0.0362 | 20.2 | < 2e-16 *** |
| *rain.wkmn* | 0.005194 | 0.00189 | 2.75 | 0.00621 ** |

Residual standard error: 0.108 on 410 degrees of freedom
Multiple R<sup>2</sup>: 0.635. Adjusted R<sup>2</sup>: 0.633
F-statistic: 238.1 on 3 and 410 DF; p-value: < 2.2e-16. AIC = -1840.2

### Analysis of factors affecting egg length in *G. pallidipes* and *G. m. morsitans Individual flies*

Correlation matrices for egg length versus six environmental variables, plus maternal ovarian category and wing length, showed that egg length in both species was highly correlated with a variety of meteorological factors. These included temperature, relative humidity, saturation deficit, NDVI, rainfall and the number of hours per day suitable for feeding activity, all averaged over the estimated period of time when the egg had been developing as an oocyte in its mother’s ovary (**Supporting Information S10**, Table S10.1 A, B). For both species, egg length was significantly reduced for *c* = 1 relative to older flies, in accord with the results in Figure 7C, D. We could detect, however, neither a progressive increase in egg length with maternal age, nor any indication that egg size decreases in the oldest flies.

There were strong correlations between various meteorological variables: NDVI, relative humidity and rainfall all provided measures of moisture; saturation deficit provided a combined measure of heat and dryness; and the number of hours suitable for tsetse feeding behaviour each day is a function of temperature. These high correlations would confuse any attempt at multiple regression modelling involving all candidate explanatory variables. Moreover, full-model regressions of egg length for individual flies accounted for only 18% of the variance for *G. pallidipes* and 21% for *G. m. morsitans* (**Supporting Information S10**, **Table S10.2 A, B**). We therefore carried out all further regression modelling using mean lengths of eggs, and of corresponding environmental predictor variables, averaged weekly for *G. pallidipes* and monthly for *G. m. morsitans*.

### Flies pooled over weeks/months

Pooling was based on the estimated date when oogenesis was completed. Weekly averaging provided 525 weeks of *G. pallidipes* averages, for the full period 1988-1999, 479 of which included five or more eggs in their averaging, weeks with fewer than 5 eggs having been excluded from the analysis. Models were developed using 341 of these weeks which lay in the period starting in week 47 (November 1991) – the first week for which weekly logger data were available. Regression using all candidate predictors accounted for 77% of the variance (**Supporting Information S10, Table S10.3 A**). Note the high *t*-value for mean maternal wing length, despite it being averaged over many flies.

Using the analogous approach for monthly-pooled averages for *G. m. morsitans* data provided 124 months for the full period (1988-99), 100 of which were based on a minimum of five eggs: models were based on 65 of these months, starting in December 1991 when the logger data became available. Regression using all candidate predictors accounted for 66% of the variance and, as with *G. pallidipes*, maternal wing length (*wlmm*) had the highest *t*-value (**Supporting Information S10, Table S10.3 B**).

To make true predictions of egg length, for 1988-91, we did not include mean *wlmm*, from our sampled tsetse data from that period, as a predictor variable; if we use tsetse data from that period, then predicting egg length is moot - we could simply obtain egg lengths directly from the data. Instead, we set aside the observed egg lengths from 1988-91 as a check on values from our true predictions. Thus, only the three variables *Tbar*, *ndvi*, and *rain* are available for driving predictions during 1988-91. Stepwise regression was run, based on the AIC criterion, to select the best model from the above three candidates and showed that, for both species, mean rainfall could be dropped with no significant change in AIC. Accordingly, the models used for prediction of egg lengths in the period 1988 – 1991 are based only on mean temperature and NDVI. Results of the final regression models are shown in Table 5A, B and were used in the construction of Figures 8 and 9. For *G. pallidipes*, the simpler regression accounted for 69% of the variance in egg length, less than that for the full model including maternal wing length and additional environmental factors. Nonetheless, the model produced a good fit both to the data collected after November 1991 (Fig. 8A), and it also provided true predictions for 1988-91 that matched well the observed mean egg lengths from that period (Fig. 8B).

Given the much smaller sample sizes for *G. m. morsitans*, the variability in the observed pooled egg lengths was much greater than for *G. pallidipes*. Nonetheless, there is good correspondence between the observed and predicted egg lengths, in both the model fitting and model prediction periods (Fig. 9). For both species, minimum egg lengths were closely aligned with peaks in the mean temperature, increasing rapidly thereafter as NDVI values increased (Figs. 8, 9 C and D).

### Analysis of factors affecting wing length in *G. pallidipes* and *G. m. morsitans* **A.** G. pallidipes

#### Individual flies

We modelled the effect of mean meteorological conditions, prevailing at the time a female tsetse was developing in its mother’s abdomen, on the wing length of the resulting adult. Product-moment correlations showed that mean saturation deficit, relative humidity, and NDVI were equally strongly correlated with wing lengths. However, linear regression of length on all 6 mean meteorological variables explained only 24% of the variance in wing lengths of the 48,795 individual *G. pallidipes* (**Supporting Information S11. Table S11.1**). Accordingly, further regression modelling used only the averages from the weekly pooled groups of flies.

#### Weekly pooling

For *G. pallidipes* all further analyses involved data pooled over weeks. The matrix of correlations between mean wing length and six predictor variables showed that there were strong correlations between mean wing length, saturation deficit, relative humidity, NDVI and the number of hours available for feeding (**Supporting Information S11. Table S11.2**). For the pooled data, linear regression, using all six available independent variables, accounted for 71% of the variance in wing length (**Supporting Information S11. Table S11.3).** Note that temperature (*Tbar*) and relative humidity (*RHbar*) are high correlated with saturation deficit *(SDbar*), and removing the first two of these variables from the regression did not affect the R^2^ value. The full-model regression could not, however, be used to predict wing lengths prior to November 1991 – since saturation deficit and the numbers of hours available for feeding (*HrsFeed*) were not measured during that period. Accordingly, we modelled wing length as a function only of temperature, NDVI and rain. This regression accounted for 64% of the variance in wing length (Table 6). Stepwise selection based on AIC showed that all three of these variables contributed predictive value.

#### Observed and predicted wing length November 1991 – December 1999

A plot of observed mean weekly wing length, overlaid with model predictions driven by mean temperature, NDVI and rainfall, showed good agreement between the two time series (Fig. 9). Weekly mean wing lengths at time of capture were consistently synchronized with annual cycles in NDVI experienced during fly development (Fig. 9A, C). NDVI increased rapidly after the onset of the rains in December and declined steadily when the rains stopped in about March. Mean wing size increased rapidly with increasing NDVI during the last quarter of each year, peaking shortly after the peak NDVI in January-February. Subsequently, an annual decline in mean wing length usually continued to follow NDVI, with a lag of 1-2 months.

There was an exception in 1999, when wing length remained high throughout most of the year. Peak annual temperatures experienced by flies during their development coincided with minimum length at time of capture (Fig. 9A, C), although the annual coincidence between lower temperatures and greater size was less consistent.

#### Observed and predicted wing length September 1988 – November 1991

Having used flies captured after 16 November 1991 to develop a predictive model for wing length as a function of *Tbar, ndvi* and *rain*, we applied the predictions to the period between the start of the study in 1988 and 16 November 1991. We did no further fitting to the earlier data; we simply used the temperature, NDVI and rainfall data during that period to make true predictions of the weekly mean wing lengths. Figure 9B shows that there was good agreement with the observed values.

### B. G. m. morsitans

#### Individual flies

For this species we had full histories for 7531 individual females, an order of magnitude fewer than for *G. pallidipes*. As expected, full-model regressions of wing length against all 6 meteorological variables, carried out at the level of individual flies, had an *R*^2^ value of only 0.19 (**Supporting Information S12**, Table S12.1). Weekly averaging of flies resulted in 25% of the 387 weeks in 1991-99 having samples of 5 or fewer flies, and 50% of the weeks having 10 or fewer. Such small sample sizes would have generated noisy weekly means of wing length, with wide confidence intervals. We therefore carried out further regression modelling only on the means from monthly groups of flies,

#### Monthly pooling

Flies were assigned to monthly groups, based on their estimated larviposition date. After monthly averaging, only 5 of 92 months had fewer than 10 flies. We ran two regression models, based on monthly averages for the 1991-99 period. For the full model, using all meteorological variables showed that NDVI, *R*^2^ = 0.66 – a good result, given the extensive averaging of the data (**Supporting Information S12**, Table S12.3).

#### Observed and predicted wing length November 1991 – December 1999

We then used a model based only on *ndvi* and *Tbar* (Table 9) – variables that were also used for the final *G. pallidipes* model. For this simpler model, *R*^2^ was 0.57, slightly less than for the full model. Based on model AIC, mean rainfall did not add any predictive value to the simpler model. Both predictors were significant, with coefficient signs the same as the similar model for *G. pallidipes*. NDVI and temperature were thus the primary drivers for both species, with qualitatively similar effects – despite the two species being modelled on different time scales. The model provided good fits to the monthly averaged wing length data for December 1991 – December 1999 (Fig. 10) and, as with *G. pallidipes*, changes in wing length were quite closely synchronized with those in NDVI.

**Figure 10.**
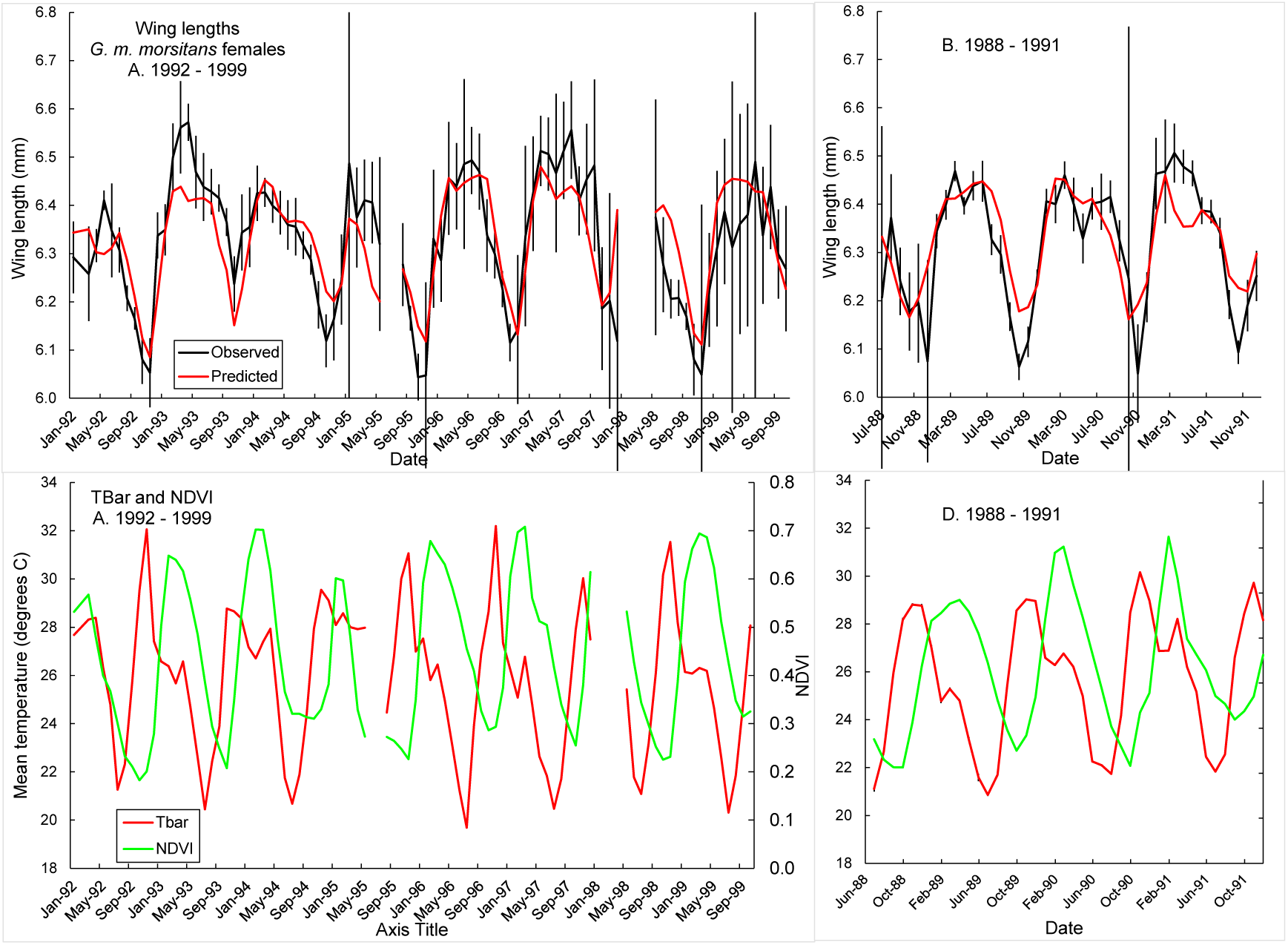
Monthly mean wing lengths of adult *G. m. morsitans* females at capture (black dots, with 95% confidence intervals), plotted against the month of their larviposition. Solid red line shows the lengths predicted by the model detailed in Table 9.

**Table 9.** Mean monthly wing lengths of female *G. m. morsitans* regressed on monthly means of mean daily temperature (*Tbar*) and *ndvi* Call: lm(= length.momn ∼ *Tbar*.momn + *ndvi*.momn, data = modl.dat)

| **Residuals:** | **Minimum** | **1Q** | **Median** | **3Q** | **Max** |
| --- | --- | --- | --- | --- | --- |
|  | -0.231 | -0.0593 | 0.00055 | 0.0427 | 0.398 |
| **Coefficients:** | **Estimate** | **Std. Error** | **t value** | **Pr(>\|t\|)** |  |
| (Intercept) | 6.54 | 8.73e-03 | 74.88 | <2e-16 *** |  |
| *Tbar.momn* | -0.0170 | 3.09e-04 | -5.496 | 3.7e-07*** |  |
| *ndvi.momn* | 0.515 | 6.10 e-04 | 8.435 | 5.5e-13 *** |  |

Residual standard error: 0.0896 on 89 degrees of freedom
Multiple R-squared: 0.569, Adjusted R-squared: 0.559
F-statistic: 58.7 on 2 and 89 DF, p-value: < 2.2e-16

#### Observed and predicted wing length November 1991 – December 1999

Following the same procedure as for the *G. pallidipes* we see that the model (Table 9) fitted to data from November 1991 – December 1999 (Fig. 10A), also provided – without further fitting – good true predictions for the 1988-91 period (Fig. 10B).

## Discussion

In producing detailed life histories of individual female tsetse, our study appears to differ from all earlier work on tsetse and on field studies on any other insect species. In the past, insect developmental rates have been used to predict the future development and, particularly, emergence dates of insect crop-pests – with the aim of optimizing the timing of insecticide application. This procedure has a venerable history (Réaumur, R.A., 1735) and there are several reviews of recent work (Damos & Savopoulou-Soultani, 2012; Rebaudo & Rabhi, 2018; Pollard et al., 2020). We reversed the procedure used in such studies: instead of predicting the future development of an insect, we assign to field-sampled female tsetse a timetable of life stages in the flies’ pasts. We work our way back from the date of capture of a female adult through: (i) the end and start dates of each pregnancy and ovulation; (ii) the date of emergence of the adult fly from its puparial case; (iii) the time underground as a larva and pupa back to the date of larviposition; (iv) the egg and larval stages in the uterus going back to the date of ovulation; (v) the time spent in the mother’s ovary as an oocyte, beginning with the start date of oogenesis.

We can therefore assess the effects of meteorological conditions, at any chosen period of a fly’s life history, on the state of the adult fly at the time of its capture. This has been achieved using a combination of published estimates of development rates in different tsetse life stages, and detailed ovarian dissection data, which facilitate age estimates with an error of only a few days (Dransfield *et al*., 1989; Rogers *et al*., 1984; Hargrove, 1994). For the first time in studies of size in tsetse, these estimates were made on females that had ovulated up to three times. Our study was thereby able to analyze an order of magnitude more flies than in previous studies with tsetse (*cf.* Table 3 and **Supporting information S13, Table S13.1**).

The advances provided a more precise picture of the factors correlating with size in tsetse, allowing us to confirm and extend some findings, and improve or correct others. We provide the first study of the correlations of meteorological and maternal factors on egg length. For both species we found that the first egg produced by an adult female was significantly shorter than all subsequent eggs, but also that there was no evidence to suggest that egg length declines in the oldest females. Similarly, several other studies in the laboratory (Langley & Clutton-Brock, 1997; McIntyre & Gooding, 1998) and the field (English et al., 2016; Hargrove et al., 2018) found no decline in the weights of pupae produced by old tsetse. These findings are at variance with a theoretical study suggesting that such a decline should occur – given 35 different scenarios defining elements of tsetse population dynamics, feeding patterns and energetics (Barreaux et al., 2022). The authors cite a single laboratory investigation in which there was indeed a decline in the weights of pupae produced by *G. m. morsitans* (Lord et al., 2021). Barreaux et al. (2022) cautioned that the flies used in that study had been in the laboratory for many generations, thus giving results that might not be valid for wild populations. However, when newly emerged tsetse were released in Zimbabwe, there was no significant difference between the survival and behavior of flies emerging from field-collected, and laboratory-produced, pupae (Vale et al., 1976). Further work is necessary to resolve this issue.

Both egg and wing lengths showed strong annual cycles, in concert with key environmental variables. At the individual level, however, the environmental and biological variables we measured accounted for a disappointingly low percentage of the variance in fly size. We are thus unable to suggest the factors that control the large, unexplained variability (∼80%) in the egg and wing sizes of individual flies. It is unclear whether it is possible for uncontrolled field studies, such as ours, to see the effects of such factors. These could be related to genetic, microhabitat or microclimate factors that we were unable to investigate.

### Predicting future changes in egg and wing lengths

Much better fits to egg and wing length data were obtained when we used weekly or monthly averages of fly sizes and regressed these lengths against time-averaged meteorological conditions prevalent during the periods of oogenesis and pregnancy. Crucially, we then showed that the fitted equations, based only on measured values of temperature and NDVI, provided good predictions of egg and wing lengths collected at other periods – without any further fitting to those data. These results suggest that we can use the fitted equation to predict how fly size will change in the future, thereby providing interesting information on the expected effects of changing climate on tsetse biology. Temperature increases and humidity decreases, in areas like Rekomitjie, will result both in increased adult mortality and reduced size of pupae with consequent increased mortality among pupae and, particularly, newly emerged adults. Simultaneously, however, similar changes in areas that are currently too cool for tsetse populations to thrive could result in meteorological patterns that are more suitable for the fly.

### What drives size changes in tsetse?

No convincing causal link has previously been established between annual cycles in fly size and meteorological variables. While wing length was correlated with saturation deficit (Jackson, 1953; Bursell & Glasgow, 1960) further data weakened rather than strengthened the correlation (Glasgow & Bursell, 1961). Similarly, while low tsetse population densities are correlated with high rates of evaporation (Nash, 1933; Jack, 1942; Jackson, 1948b) tsetse are remarkably resilient to dehydration, both as pupae and as mature adults (Bursell, 1958, 1959).

By contrast, for populations of *G. m. morsitans* and *G. pallidipes*, of both sexes, on Antelope Island, Lake Kariba, Zimbabwe, mortality among post-teneral adults was always more highly correlated with temperature than with relative humidity, or saturation deficit (Hargrove, 2001a). Given that, every 2-3 days, adults take large bloodmeals consisting of 81% water, by weight (Taylor, 1976), this is unsurprising. The water balance issue for non-teneral adults is not shortage of water but, rather, the need for rapid elimination of water after each meal.

Teneral flies, which have a softer and more permeable exoskeleton for some hours after emergence and are more prone to desiccation at low relative humidity, have often been identified as the weakest link in the chain of tsetse life. Bursell (1959) provides evidence, however, that – even for tenerals – low water content is not the critical issue; instead teneral mortality was more likely due to flies having insufficient fat to stay alive until they could find their first bloodmeal.

Rogers pioneered the use of satellite imagery in the study of tsetse and produced convincing evidence that tsetse distribution, population density, adult mortality, adult size, as well as seasonal changes in numbers of human trypanosomiasis, are all highly correlated with NDVI (Rogers, 1991, 2000; Rogers & Williams, 1993; Rogers et al., 1996). As with earlier authors, however, he did not establish a causal link showing how NDVI might act on tsetse to affect the above-mentioned features of tsetse and trypanosomiasis biology. The present study is no exception in finding that measures of wetness – such as relative humidity, saturation deficit and NDVI – provide the highest correlations with egg and wing lengths. We are still faced, however, with the same problem as earlier workers: the need to establish a causal chain between such measures and fly size and other features of tsetse biology (Glasgow & Bursell, 1961). We cannot claim to have solved the problem – but we advance below alternative interpretations of the available data.

### It’s all about the food?

We suggest that attempts to establish a causal link, via direct effects of dryness on tsetse failed, because no so such direct effects exist. Thus, whereas NDVI and relative humidity are demonstrably highly correlated with various measures of tsetse population health, there seems no reason to think that they have, or could have, any direct effect on tsetse (Bursell, 1959). Indeed, NDVI could take values that are optimal for tsetse in a given area, and tsetse would still be entirely absent if, for whatever reason, there were no vertebrate hosts on which tsetse could feed. Indeed, complete absence of herbivores would favor an increase in vegetative growth and thus NDVI and, simultaneously, ensure the complete absence of tsetse.

Conversely, Antelope Island, Lake Kariba, Zimbabwe, provides an example of an arid area where populations of tsetse thrived. The island is a 5 km^2^ rocky outcrop, formed when the Zambezi River was dammed. The vegetation consists of sparse grass and deciduous trees. For a large portion of the year there is thus high saturation deficit, low relative humidity, minimal photosynthetic activity and NDVI. The island thus appears entirely unsuitable for tsetse. Yet populations of *G. m. morsitans* and *G. pallidipes* grew on the island at rates seldom equaled elsewhere – for the simple reason that the island was stocked with 33 head of cattle, fed on hay and concentrate, and paddocked to ensure a constant source of hosts, readily available to the tsetse (Vale et al., 1986).

Consideration of the above artificial situations suggest ways in which NDVI and relative humidity could, indirectly, affect the size, and general health of natural tsetse populations. In most such natural situations, we expect a positive correlation between NDVI and local densities of the vertebrates (wild or domestic) that are the sole source of food for both sexes of every species of *Glossina*. If declining NDVI is correlated with declining vertebrate densities, it will become more difficult for female tsetse to locate a host and to obtain the regular bloodmeals required to nourish themselves and their offspring.

Annual cycles at Rekomitjie provide a good illustration of the above idea. The rapid decline in NDVI, following the onset of the dry season at Rekomitjie, reflects not only reductions in growth of green plants – but also the rapid consumption of that green vegetation by the herbivores that are also the hosts for tsetse. As the vegetation disappears, hosts may be expected to move in search, literally, of greener pastures. The probability of a tsetse encountering a host then decline. The importance of host density for the well-being of a tsetse population was illustrated by a game destruction exercise carried out between 1962 and 1967 in an enclosed area of 541 km^2^ near Nagupande in western Zimbabwe. Destruction of almost all (>2000) of the mammals in the area led, unsurprisingly, to the almost entire elimination of the *G. m. morsitans* population – which seemed only to be sustained by a small amount of immigration (Lord et al, 2017). More importantly, a 50% reduction in the game population led to a 95% reduction in the numbers of flies caught on fly rounds – in keeping with the increased difficulty tsetse experience in feeding as host density declines.

The effects of cyclical changes in local host density at Rekomitjie are compounded by the effects of temperature. As temperatures increase, between July and November, tsetse face the problems that, being poikilotherms, they use energy more rapidly (Taylor, 1978 a, b; Hargrove & Coates, 1990) – and need to feed more frequently. Simultaneously, however, the number of hours per day suitable for tsetse feeding activity declines sharply (Fig. 2A). The problem is compounded for females and their offspring. During a 9-day pregnancy females feed every 2.5 – 3 days (Randolph et al., 1992; Hargrove, 1999a, b; Hargrove & Williams, 1999) and therefore take at least three blood meals. As the pregnancy duration decreases with increasing temperature the female would thus still need at least three blood meals to produce a full-sized larva – but with fewer days, and fewer hours per day, in which to find those meals. These factors, together with declining local host densities, combine to make it increasingly difficult for females to feed sufficiently frequently to build up sufficient reserves to produce the size of larva that are possible at cooler, moister times of the year. An indirect effect of temperature on adult mortality is, similarly, consistent with the increased number of hunger-related deaths resulting from the increase in the feeding rate with increasing temperature (Hargrove, 2001b).

The observed decline with temperature in the lengths of eggs, and of the wings the resulting adults, are thus to be expected. Declining fly size brings a further problem, however, because there is severe selection at the hottest times of the year against the smallest tsetse, particularly because they also have relatively less fat than larger flies (Phelps & Clarke, 1974). When the rains start, in December, temperatures decrease, humidity and NDVI levels increase, rates of vegetative growth are greatly accelerated, and many host species give birth in the following months. The previously declining trends in fly size are thereby all reversed; females produce larger eggs and larvae and, thereby, offspring often larger than their mothers (Fig. 6A, B).

Moreover, the rate of increase in the mean fly size accelerates because, between October and January, small flies are strongly selected against (Fig. 8). Contrary to the findings of earlier studies, we show that this selection extends for some weeks after the teneral period. The larger surviving flies produce larger pupae (Table 9) and the larger adult daughters produce larger eggs (Table 5). The regular annual cycle in egg lengths may then be seen, largely, as a secondary result of changes in wing length: egg length is positively correlated with maternal size, though it is also a function of the conditions the adult female experiences during oogenesis.

### Bigger is not always better

Most previous studies have focused on the loss of small tsetse at very early stages of adult life. Only Glasgow (1961) has adduced evidence suggesting that there should also be selection against larger flies, as must be the case: thus, whereas there are clearly physical constraints to the size of larva a female can produce, if each female produced an offspring slightly larger than herself, size would increase without limit. Instead, what we expect, as found by Galton (1886) for human males, is that there will be a “regression towards mediocrity”; very small females will tend to produce daughters larger than themselves, and very large females to produce daughters that are smaller.

### Limitations of our study

#### Sources of error

The large proportion of variation unaccounted for by regression of egg and wing lengths of individual flies, on meteorological variables, was due to a variety of factors: (i) Measurement error: regression analysis demonstrated significant differences between egg length measurements made by five dissectors using eight different microscopes (**Supporting Information S14**, Table S14.1). (ii) Fly size exhibits remarkably small absolute variation: the difference between maximum and minimum monthly mean length values is only *c*. 0.5 mm and *c*. 0.2 mm for wing and egg lengths, respectively (**Supporting Information S15**, Figure S15.1). Microscope measurements, made even at the highest magnification, only allowed lengths to be estimated to the nearest 0.02 mm, *i*.*e*., about 10% of the difference between maximum and minimum egg length, even without variations between dissectors and microscopes. (iii) We assumed no uncertainty in the rates of development used in generating life histories and assumed no effect of fly size and fly location on these rates. The models, applied to the Rekomitjie situation, were model fits to data, sometimes from laboratory flies. Moreover, in the absence of measured rates for *G. pallidipes* we used rates derived for *G. m. morsitans*. While temperature, the driver of the rates, is accurately measured by a weather station, the fly will almost always be experiencing a different temperature (Vale, 1971; Torr & Hargrove, 1999). Violations of these assumptions, along with age estimation errors, may well lead to errors of a few to several days, in the timing of egg and wing developmental periods. However, wetness, greenness and temperature change slowly on a longer timescale. This may explain why we were so much more successful with modelling of weekly and monthly pooled data.

Whereas averaging the attributes of many flies over weeks or months provided useful predictive models, the disadvantage is that maternal attributes, such as size and ovarian age, can no longer be linked to individual daughters, thus inhibiting the understanding of maternal effects. Thus, while it seems certain that maternal size and age affect offspring size, and we see such effects from regressions carried out on individual flies, we cannot test these ideas on pooled samples.

### Difficulties in estimating chronological ages

Carey et al (2012) point out that technologies for estimating the age of individual insects are generally accurate for young flies, less so for mid-aged individuals and that none can differentiate ages within the oldest age groups. Tsetse provide something of an exception in this regard: in this study we have assigned ages to flies up to *ca*. 35 days old – by which age we may expect that >50% of female tsetse will have died, even in favorable field conditions (Hargrove *et al*., 2011). There is still, however, no way of pinpointing the exact number of ovulations completed by all female tsetse for *c* = 4+4*n* to 7+4*n* (Hargrove, 2020) and we were thus unable to assign an age to 20-30% of the older dissected flies. Since wing and egg lengths varied little for flies where *c* >2 (Fig. 7), minimal bias should result from the loss of the older flies. Nonetheless, the inability to estimate accurately the chronological ages of older flies is undeniably a shortcoming of the ovarian dissection approach.

The problem has been cited in a paper suggesting that the combined use of mid-infrared spectroscopy and machine learning, could be implemented as a routine surveillance tool, in the control of vectors such as tsetse, for the accurate determination of sex and chronological age (Pazmiño-Betancourth *et al*., 2024). It is suggested further that the method could thus replace the use of ovarian dissection, which is viewed as labor-intensive, slow and inaccurate. Whereas, however, it is claimed that artificial intelligence could be used to discern the sex of a tsetse fly with 96% accuracy – natural intelligence and a cursory glance at the tip of the ventral abdomen would make the gender distinction with 100% accuracy. Moreover, the claimed ability of the technique to be able to separate female flies of ages 3 days and 5 weeks with 87% accuracy is scarcely impressive. A trained dissector would, again, make the same distinction with close to 100% accuracy.

More importantly, the authors are missing the opportunity afforded by their new approach, which would come from using it in concert with, rather than replacing, ovarian dissection. Thus, whereas ovarian dissection cannot be used, by itself, to determine the number of times (*c*) a fly has ovulated if *c* > 3 it can identify this number modulo 4. For example, we do know that a fly where *c* = 4+4*n* has ovulated exactly 4, 8, 12, 16 *etc*. times. At the start of these ovulation cycles a female, living at a mean temperature of 25°C, would be order 34, 71, 107, 143 days old, respectively. If, therefore, infrared technology could accurately separate any two females, differing in age by order 36 days, it could be used – together with ovarian dissection – to provide highly accurate estimates of the chronological ages of female tsetse of all ages. One would then be able to produce the full life histories for all these flies – which would be a remarkable analytical breakthrough. We caution, however, that the same breakthrough was promised with the combined use of ovarian dissection and pteridine analysis (Lehane & Mail, 1985; Lehane & Hargrove, 1988) but the method has never been developed. The infra-red study must be repeated, using field flies of known chronological age, so that its true usefulness can be assessed.

### Relevance of our study

Knowledge of the chronological ages of individual insects sampled in the field is pivotal in a host of areas associated with the study and control of insect vectors of disease. These include population dynamics and ecology, including age-dependent estimates of mortality and fecundity, age-dependent incidence and prevalence of infection and, thus, of vectorial capacity, and the assessment and improvement of vector and disease programs. Our methodology could also be used to assess and calibrate new approaches to the problem of estimating age in field-sampled insects.

Since our study appears to be unmatched in its scope by published work on tsetse, or any other fly, it serves as a model for similar work that could be carried out on other insect vectors of disease. It is now more than 75 years since Detinova (1949, 1962) published her groundbreaking study on the ovarian dissection of mosquitoes, serving as a model for the development of analogous techniques for other insects – applied to tsetse by Saunders (1960) and refined by Challier (1965). But the technique has never been used successfully in field studies of mosquitoes, and defining the age of a wild-caught mosquito remains a challenging, unreliable process (Johnson et al., 2020). If, however, ovarian dissection and new technologies could be combined, as we have suggested for tsetse, it might yet be possible to determine accurately the chronological ages of individual mosquitoes.

## Conclusions

We have developed a novel methodology for mapping a complete life history of field-collected female tsetse of ages up to *ca*. 35 days. This allowed us to demonstrate the effect, on a fly’s egg and wing lengths, of the meteorological conditions experienced by her mother while the fly was developing as an oocyte in the ovaries and as an egg or larva in the uterus. We demonstrate the central importance of NDVI and the highly correlated humidity, and temperature, in determining fly size – and show that selection against under-sized individuals in the hot-dry season continues for some weeks after emergence rather than being restricted to mortality in teneral adults, as previously thought. Most importantly we propose a mechanism for causal links between tsetse size and meteorological factors, which envisages indirect effects of measures of dryness and heat on the general well-being of tsetse populations via effects on the ease and safety with which the flies can feed. The methodology has important implications for control, evolution, population dynamics.

## Acknowledgements

We thank Glyn Vale, Stephen Torr and Brian Williams for their valuable comments on the manuscript.

## Author Contributions

Conceptualization: John Hargrove

Data curation: John Hargrove

Formal analysis: John Van Sickle, John Hargrove, Faikah Bruce

Investigation: John Hargrove, John Van Sickle

Methodology: John Van Sickle, John Hargrove

Project administration: John Hargrove

Software: John Van Sickle, John Hargrove, Faikah Bruce

Supervision: John Van Sickle, John Hargrove, Faikah Bruce

Validation: John Hargrove, John Van Sickle

Visualization: John Van Sickle, John Hargrove, Faikah Bruce

Writing – original draft: John Hargrove, John Van Sickle.

Writing – review & editing: John Hargrove, John Van Sickle, Faikah Bruce

## Captions for Supporting Information

**Supporting Information File S1.** Daily meteorological readings.

**Supporting Information File S2.** Interpretation of ovarian dissections.

**Supporting Information File S3.** Steps required to generate life history of female tsetse.

**Supporting Information File S4.** Table of estimated life histories of female *G. pallidipes*.

**Supporting Information File S5.** Circannual variation in meteorological variables.

**Supporting Information File S6.** Hours per day suitable for tsetse feeding activity.

**Supporting Information File S7.** Distribution of egg and wing lengths.

**Supporting Information File S8.** Egg length vs maternal age and of month of capture: monthly pooling

**Supporting Information File S9.** Wing length *G. m. morsitans* vs ovarian category and month when the fly emerged as an adult.

**Supporting Information File S10.** Analyses of factors affecting egg lengths in individual *G. pallidipes* and *G. m. morsitans*.

**Supporting Information File S11.** Analyses of factors affecting wing lengths in female *G. pallidipes*.

**Supporting Information File S12.** Analyses of factors affecting wing lengths in female *G. m. morsitans*

**Supporting Information File S13.** Sample sizes in previous studies.

**Supporting Information File S14.** Egg lengths vs dissector and microscopes.

**Supporting Information File S15.** Wing and egg lengths emphasising small relative change.

